# Replenishing the malaria drug discovery pipeline: Screening and hit evaluation of the MMV Hit Generation Library 1 (HGL1) against asexual blood stage *Plasmodium falciparum*, using a nano luciferase reporter read-out

**DOI:** 10.1101/2022.06.07.495126

**Authors:** Koen J. Dechering, Martijn Timmerman, Kim Rensen, Karin M.J. Koolen, Saman Honarnejad, Martijn W. Vos, Tonnie Huijs, Rob W.M. Henderson, Elodie Chenu, Benoît Laleu, Bailey C. Montefiore, Matthew D. Segall, James E. J. Mills, Eric M. Guantai, James Duffy, Maëlle Duffey

**Affiliations:** TropIQ Health Sciences, Transistorweg 5, 6534 AT Nijmegen, The Netherlands; Pivot Park Screening Centre, Oss, North Brabant, The Netherlands; Medicines for Malaria Venture, Route de Pré-Bois 20, PO Box 1826, 1215 Geneva 15, Switzerland; Optibrium, F5-6 Blenheim House, Cambridge Innovation Park, Denny End Road, Cambridge CB25 9PB, United Kingdom; Sandexis Medicinal Chemistry Ltd, Innovation House, Discovery Park, Sandwich, CT13 9FF, United Kingdom; Department of Pharmacy, Faculty of Health Sciences, University of Nairobi, 00202-Nairobi, Kenya

**Keywords:** Malaria, high throughput screening, phenotypic screen, drug discovery, nanoluciferase, screening cascade

## Abstract

A central challenge of antimalarial therapy is the emergence of resistance to the components of artemisinin-based combination therapies (ACTs) and the urgent need for new drugs acting through novel mechanism of action. Over the last decade, compounds identified in phenotypic high throughput screens (HTS) have provided the starting point for six candidate drugs currently in the Medicines for Malaria Venture (MMV) clinical development portfolio. However, the published screening data which provided much of the new chemical matter for malaria drug discovery projects have been extensively mined. Here we present a new screening and selection cascade for generation of hit compounds active against the blood stage of *Plasmodium falciparum*. In addition, we validate our approach by testing a library of 141,786 compounds not reported earlier as being tested against malaria. The Hit Generation Library 1 (HGL1) was designed to maximise the chemical diversity and novelty of compounds with physicochemical properties associated with potential for further development. A robust HTS cascade containing orthogonal efficacy and cytotoxicity assays, including a newly developed and validated nanoluciferase-based assay was used to profile the compounds. 75 compounds (Screening Active hit rate of 0.05%) were identified meeting our stringent selection criteria of potency in drug sensitive (NF54) and drug resistant (Dd2) parasite strains (IC_50_ ≤ 2 µM), rapid speed of action and cell viability in HepG2 cells (IC_50_ ≥ 10 µM). Following further profiling, 33 compounds were identified that meet the MMV Confirmed Active profile and are high quality starting points for new antimalarial drug discovery projects.

## Background/introduction

Along with AIDS and tuberculosis, malaria remains one of the deadliest human pathogen-borne diseases, accounting for 241 million cases and 627’000 estimated deaths in 2020 ^1^. According to the World Health Organization (WHO), nearly half of the world’s population is at risk, and malaria disproportionally affects children under the age of five and pregnant women in Sub-Saharan Africa ^2^. Out of the nearly 200 species included in the *Plasmodium* genus, five of them are known to induce malaria in humans: *P. falciparum*, *P. vivax*, *P. ovale*, *P. malariae* and *P. knowlesi*. *P. falciparum* infections represent up to 90% of the worldwide burden, followed by far by *P. vivax* (3%) ^2^, and is the only species easily adapted to *in vitro* culture ^3^. The *P. falciparum* life cycle, a complex interplay between female *Anopheles* mosquitoes and human hosts ^4^, allows various intervention points for treatment, prevention and transmission blocking, all of which are essential to the eradication of the disease. Although tremendous progress has been achieved regarding drug-based malaria control over the last two decades, mainly due to the introduction of Artemisinin Combination Therapies (ACTs), significant challenges remain to reach this goal. One major hurdle is the emergence of genome alterations that cause resistance. In particular, a resistance phenotype to artemisinin and its derivatives used in ACTs, characterized by an extended clearance time of the parasites from the treated patients, has been emerging in South-East Asia in the late 2000’s ^5^ and is now spreading to Africa ^6^, potentially compromising the proper efficacy of the ACTs. This emphasizes the urgent need for new antimalarial drugs with alternative modes of action and mechanisms of resistance. Phenotypic screens, or whole cell-based assays, have successfully been used in the past to identify such new classes of compounds, leading for instance to the current antimalarial clinical candidates Cipargamin (KAE609) ^7^ and MMV533 ^8^. In contrast with target-based screening and drug design approaches that focus on single targets, target-agnostic phenotypic screening is a powerful complementary hit discovery approach that exploits cellular and organism-level physiology to unbiasedly identify compounds with new modes of action and potential polypharmacology. Major improvements in robotics and flexible automation over the last decades have made High-Throughput Screening (HTS) a mainstay in hit discovery, allowing rapid exploration of large drug-like compound libraries covering a broad chemical space.

The first major whole cell phenotypic antimalarial HTS campaigns were described by the Genomics Institute of the Novartis Research Foundation (GNF) in 2008 ^9^and then by GSK ^10^ and St. Jude Children’s Research Hospital (SJCRH) ^7^ in 2010. The last large HTS was reported by Griffiths University and MMV in 2014 ^11^. The chemical matter from these HTS, the majority of which has been made public in ChEMBL, has been extensively investigated and the opportunity to use these data sets for the identification of novel starting points for a drug discovery project is exhausted. Lessons can be learnt from previous screening campaigns and the knowledge can be used to increase the likelihood of discovering new chemical series with potent antimalarial activity. The identification of new high quality, drug-like, chemical series is dependent on the quality and diversity of the compounds tested. Malaria drug discovery projects cannot afford the luxury of testing compounds with inherent developmental challenges based on either a lack of structural novelty or chemical groups associated with poor pharmacokinetics or safety. With this in mind, we have developed compound selection criteria for improving the diversity and property profile of screening libraries. We describe the Medicine for Malaria Venture (MMV)’s *P. falciparum* asexual blood stage phenotypic screening platform, exemplified with the 141,786 compounds Hit Generation Library 1 (HGL1). In addition to the publication of the complete HTS data set, new screening methodologies have been developed. A new *P. falciparum* strain genetically modified to express a nanoluciferase reporter gene driven by the *ef1α* promoter was used for the primary screen. This reporter provided a robust read-out for parasite replication with incubation times as short as 12 hours. Active molecules were subsequently validated in orthogonal assays for specificity and speed of kill, resulting in a prioritized list of 33 compounds meeting the MMV Confirmed Active criteria that can serve as starting point for new antimalarial drug discovery programs.

## Methods

### Library design

The Medicines for Malaria Venture (MMV) Hit Generation Library 1 (HGL1) was designed as a set of approximately 140,000 diverse, novel, and high-quality commercially available compounds. A stepwise approach was used to select the compounds in the library. Firstly, a set of approximately 8,700,000 commercially available compounds was created from publicly available vendor databases. The set of compounds was filtered down to 5,900,000 compounds by removing compounds containing features known to be enriched in compounds exhibiting toxicity and/or assay interference ^12^.These compounds were clustered using an in-house program to give a structurally diverse, smaller, selection of 875,000 compounds. Instead of applying strict property-based cutoff criteria, a multi parameter optimization (MPO) method was used to further refine the selection and enrich the compound collection with high-solubility, ‘drug like’ compounds. The novelty was ensured by excluding repeats of compounds in the MMV in-house database or the ChEMBL-NTD archive and that the number of compounds per cluster (including MMV compounds) did not exceed 8. The majority of the compounds selected achieved a balanced profile, but a smaller unbiased random selection was included to increase structural diversity whilst maintaining the overall quality of the library. The property profile of the library calculated using StarDrop^TM^ is shown in Figure 1, detailed in Table S1 and the calculation of the MPO score is portrayed in Figure S1.

**Figure 1:**
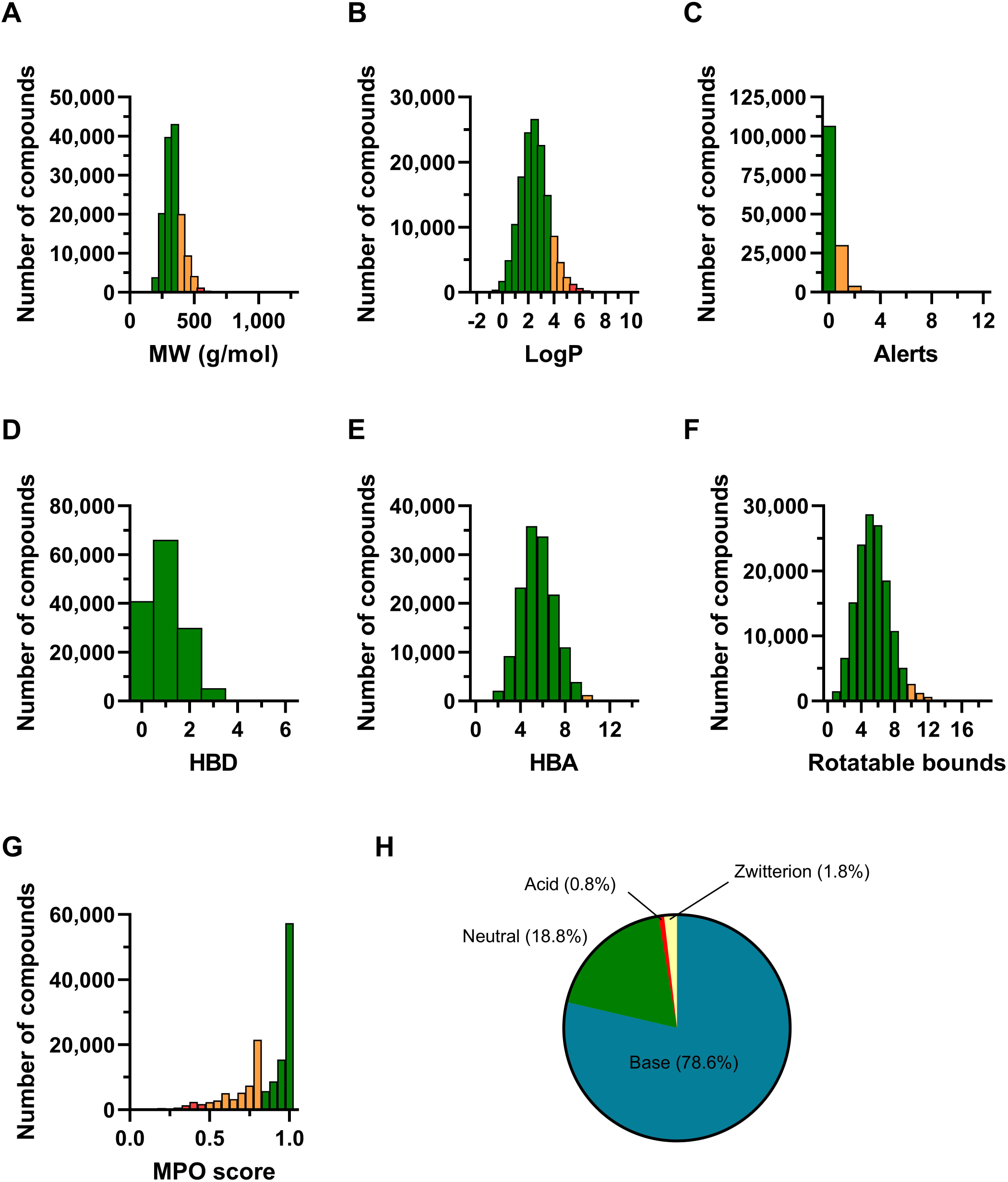
MMV Hit Generation Library 1 (HGL1) designed by promoting diversity, novelty and quality. Composition of the library in terms of molecular weight (MW, A), partition coefficient (LogP, B), structural alert (C), number of hydrogen bond donors (D) and acceptors (E), rotatable bonds (F), multiparameter optimization (MPO) score (G) and molecular class (H). These properties were calculated using StarDrop. Ideal values for each criterion are depicted in green, acceptable ones in orange, and non-ideal ones in red.

### Development of the nanoGlo assay

#### Generation of a transgenic reporter parasite

A nanoluc reporter gene under control of an *ef1α* promoter was introduced in the *pfs47* locus of *P. falciparum* NF54 parasites by gene targeting. The targeting construct was originally designed for CRISPR/Cas9 mediated integration (Figure 3A) but used here for episomal expression followed by selection for single cross-over events. For construction of the targeting plasmid the construct PLF0047 described earlier ^13^ was modified to introduce a *BsiWI* restriction site between the *pfs47* 5’ and 3’ target regions. To this end, a synthetic double stranded oligomer derived through hybridisation of oligonucleotides MWV539 and MWV540 (Table S2) was inserted in the *SalI* and *SacII* sites of PLF0047 using restriction digestion and ligation. Simultaneously a reporter gene subcloning vector was constructed as follows: the promoter region of *ef1α* (PF13_0304) was amplified from genomic DNA using primers MWV524 and MWV525 (Table S2) and introduced upstream of the nanoluciferase gene in the commercially available vector pNL1.1 (Promega) by *XhoI/NcoI* restriction digestion and ligation, followed by the introduction of the 3’ *hsp86* termination fragment from plasmid MV163 ^14^ by *XbaI/HindIII* restriction digestion and ligation. Next the complete reporter gene cassette 5’ *ef1α*-*nluc*-3’*hsp86* obtained after *BsiWI/XhoI* digestion was ligated in the modified PLF0047 construct after *BsiWI/SalI* restriction digestion, yielding the final targeting plasmid pMVAB229. The resulting targeting construct was introduced in *P. falciparum* NF54 parasites by electroporation as described previously ^14^. Parasites were placed in a semi-automated culture system ^15^ in culture medium (RPMI1640 medium supplemented with 367 uM hypoxanthine, 25 mM HEPES, 25 mM sodium bicarbonate) with 5% human type O red blood cells and 10% human type A serum. 24 hours after transfection, parasites were cultured in the presence of 2.6 nM WR99210 until viable parasites appeared upon inspection of Giemsa-stained thin smears. Episomal expression of the nanoluc reporter gene was confirmed in luminescence assays using NanoGlo substrate according to instructions of the supplier (Promega, Leiden, the Netherlands). Subsequently, episomal plasmids were cured by culturing in the absence of drug for 3 weeks. Then, cultures were again placed on WR99210 selection medium to select for parasites that had integrated the plasmid via cross-over at the targeted locus. Luciferase positive parasites were cloned by limited dilution in 96 well plates. Positive clones were transferred to the semi-automated culture system and genome integration of the reporter was verified by PCR using primers MWV300, MWV301, MWV531 and MWV537 (Figure 3A, Figure S2 and Table S2) and luminescence assays. A clone (clone f10) with single cross-over integration at the 3’ target site was selected for establishment of a high throughput parasite replication assay.

#### Luminescent asexual blood stage replication assays

An asexual blood stage replication assay by means of monitoring of nanoluciferase activity was established by seeding parasites in 384-well OptiPlates (Perkin Elmer, PN 6007299) in culture medium using a Multidrop dispenser (ThermoFisher Scientific). To establish optimal conditions, parasitemia was varied between 0.2% and 1.6% at 1%, 1.5% or 2% hematocrit. Half of the wells were treated with 10 uM dihydroartemisin and the other half with vehicle control (0.1% DMSO). Plates were incubated for 72 hours before nanoGlo substrate (Promega; N1120) was added. Luminescence signal was read in a Spectramax i3x plate reader (Molecular Devices). To increase the throughput, the 384-well plate assay setup was scaled up to a 1536-well plate format in which the incubation time, plate edge effects, readout stability and DMSO tolerance was assessed and optimized. Using the optimal parasitemia and hematocrit concentrations the parasites were seeded in 1536-well plates (Greiner, Cat#782080) using a Certus dispenser (Fritz Gyger AG). 10 µM dihydroartemisinin (reference compound) or DMSO (0.2%) was dispensed using an Echo acoustic dispenser (Beckman Coulter). The assay was incubated for 48 or 72 hours at 37 °C, 5% CO_2_. After incubation, plates were equilibrated at room temperature for 1 hour. The nanoGlo or pLDH detection reagents were added using a Certus dispenser (Fritz Gyger AG). The nanoGlo detection reagents were prepared according to the manufacturer recommendations. For pLDH detection reagents, 286mM sodium-L-Lactate (VWR; A1004.0050), 286 µM ADAP (Sigma-Aldrich; A5251), 357.5 µM Resazurin (Sigma-Aldrich; 199303) and 5.66 U/ml Diaphorase (Worthington; LS004327) were dissolved in a 0.2 M Tris-HCl solution (pH 8.0), containing 1.4% Tween-20 and 0.1% TritonX-100 (Sigma-Aldrich; 252859; P2287 and T9284). The measurements were carried out using the PHERAstar multimode reader (BMG LABTECH). The luminescence signal was measured for 0.1 sec/well and the fluorescence intensity (ex: 340 nm; em: 590 nm) of the pLDH assay was measured every 10 minutes for 90 minutes to determine the readout stability. Using the optimal concentrations and incubation time, the robustness set compound collection ^16^ was tested to assess pLDH and nanoGlo assay labilities towards various classes of assay interfering compounds. Parasites were dispensed using a Certus dispenser (5 µl/well), compounds/controls (2.5 or 10 nl) were added with the Echo. Parasites were incubated for 48 or 72 hours at 37 °C, 5% CO_2_. After incubation, the plates were allowed to equilibrate to room temperature for 1 hour. Detection reagents were added using the Certus (1.25 µl for nanoGlo or 2.5 µl for pLDH detection reagents). Immediately after the addition of the nanoGlo reagents, the assay plates were measured using the PHERAstar. After addition of the pLDH detection reagent, the plates were incubated for 45 minutes at room temperature. After incubation, the fluorescence intensity signal was measured using the PHERAstar reader (Ex 340, Em 590 nm). Analysis of the robustness set was carried out using ActivityBase XE (IDBS) and Vortex (Dotmatics) software packages. Because of evaporation effects in the outer wells, these were removed from the analysis. The signal from the remaining wells was analysed and the activity of the compounds was scaled to control conditions, using 10 µM dihydroartemisinin and DMSO as 100 and 0 % effect, respectively.

### Primary phenotypic screening and single point active confirmation

The HGL1 library of 4- and 10-mM solutions in Labcyte Echo compatible plates (LP-0400) were transferred to Pivot Park Screening Centre (PPSC) by MMV. The screen was performed using the 4 mM source plates. For the screening a picklist was used to transfer a maximum of 1176 compounds according to the assay plate layout, while columns 1-4 were left empty for controls. Compounds and controls (10 µM DHA or DMSO) were transferred to the assay plate using an Echo acoustic dispenser (2.5 nl/well). Parasites were added to each well by dispensing 5 µl parasite solution (1.2% parasitemia, 1.5% RBCs) using the Certus. Plates were placed interleaved in an incubator for 72 hours at 37 °C and 5% CO_2_. After 72 hours incubation the plates were allowed to equilibrate to room temperature for 1 hour before addition of 1.25 µl of nanoGlo reagents to all wells using the Certus. The luminescence signal was measured immediately after the addition on the PHERAstar reader. Compound effects were normalized and determined using 10 µM DHA and DMSO as maximum and minimum effect controls. All plates were analysed using ActivityBase (IDBS) and Vortex (Dotmatics) software. Compound activity was expressed as % effect and Z-score, calculated per compound.

Compounds selected for activity confirmation were cherry picked using an Echo acoustic dispenser and retested in the nanoGlo assay using the same procedure as described above. Next to the nanoGlo assay, the same compounds were retested in the pLDH assay. The same procedure as for the nanoGlo assay was used, except for the addition of the detection reagents, pLDH detection reagents (2.5 µl/well) were added to each well using a Certus dispenser. Plates were incubated for 45 minutes at RT after which the fluorescence signal was measured on the PHERAstar reader (Ex 340 nm, Em 590 nm). Compound activity was expressed as % effect and Z-score, calculated per compound.

### Dose Response Assays

Dose Response Curve (DRC) assays were performed at two different industrial sites, depending on the stage of the compounds in the profiling cascade. At PPSC, the DRC assays were carried out in a 1536-well format, using the 10 mM compound source plates and Echo acoustic dispenser for transfer to assay plates. Compound concentrations in the source plates ranged from 5 mM to 156 nM, spread over 10 concentration points and ½ log dilution series. For the assay, 10 nl of the compounds in source plates were transferred to the assay plates according to the plate layout, leaving the outer two rows/columns empty. Parasite solution was added (5 µl/well) to each well using a Certus dispenser. Plates were incubated interleaved in an incubator at 37 °C with 5% CO_2_ for 12 - 72 hours. After either 12 or 72 hours, plates were equilibrated for 1 hour at room temperature. Detection reagents were added to the plates using a Certus dispenser. For the nanoGlo assays, 1.25 µl nanoGlo detection solution was added to each well and for the pLDH assays, 2.5 µl of pLDH detection reagent was added. The signal from nanoGlo assays were measured immediately after the addition of the detection reagents using the PHERAstar reader (luminescence, 0.1 sec/well). The pLDH assays were incubated for 45 minutes at room temperature and after the incubation, the fluorescence intensity was measured using the PHERAstar reader (ex 340, Em 590). Data were analysed using Activity Base (IDBS) and Vortex (Dotmatics). Activity of the compounds was expressed as % effect, normalized between the positive and negative controls. DRCs were fitted using a four-parameter fit equation. At TCG Life Sciences (TCGLS), the DRC assay was carried out in a 384-well format, using 12.5 mM stock solutions. Concentration of the dilution series in the source plate ranged from 25 µM down to 8 nM, spread over 10 points and ½ log titrations. For the assay, 2.5 µl of the compounds in source plates were transferred to the assay plates according to the plate layout, leaving the outer two rows/columns empty. Parasite solution was added (22.5 µl/well) to each well and plates were incubated interleaved in an incubator at 37 °C with 5% CO_2_ for 72 hours. After 72 hours, plates were frozen at -80°C overnight and thawed at room temperature for 5 hours. After adding the pLDH detection reagents, the plates were shaken and incubated in the dark for 20 minutes. The absorbance at 650 nm was then measured at room temperature in a plate reader (Spectramax M5, Molecular Devices). Activity of the compounds was expressed as % effect, normalized across positive and negative controls. DRCs were fitted using a four-parameter fit equation.

### HepG2 toxicity assay

*In vitro* toxicity assays were performed at two different industrial sites, depending on the stage of the compounds in the testing cascade. At PPSC, HepG2 cells (ATCC, HB-8065) were cultured using DMEM supplemented with 0.1% Pen-Strep solution (Invitrogen; 15070-063) and 10% FBS (Gibco; 10091-148). After starting the culture, the cells were at least split twice before use in the toxicity assay. For the assay, medium was removed, and cells were washed twice with PBS. After washing, trypsin solution (Gibco; 15400-054) was added to the cells. Cells were allowed to detach from the culture flask and harvested in culture medium containing serum. Cells were spun down (5 min, 1000 rpm) and the cell pellet was resuspended in culture medium. Cell density was measured using the Luna cell counter (Logos BioSystems) and cells were diluted to a concentration of 2.10^5^ cells/ml. Compounds and positive control (10 µM f.a.c of Staurosporine (Sigma-Aldrich, Cat # S5921) were added (10 nl/well) to the assay plates using the Echo dispenser. HepG2 cells (1000 cells/well; 5 µl/well) were seeded using a Certus dispenser. Plates were incubated interleaved for 72 hours at 37 °C, 5% CO_2_. After 72 hours, the plates were equilibrated to room temperature for 1 hour. CellTiter-Glo reagent was added to all wells (1.25 µl/well) according to the manufacturer (Promega, Cat# G924B). The luminescence signal was measured immediately after addition of the detection reagents on the PHERAstar reader. At TCGLS, HepG2 cells (ATCC, HB-8065) were cultured using DMEM supplemented with 0.1% Pen-Strep solution (Gibco), 1 mM Sodium Pyruvate (Gibco), 10 mM HEPES (Sigma) and 10% FBS (Invitrogen). Cells were sub-cultured in growth media (DMEM + 10% FBS) for 48 hours prior to cell plating. For the assay, cells were seeded in 384-well plates with a cell count of 2000 cells per well in 50 µl of media and incubated for 24 hours in a 37 °C, 5% CO_2_ incubator so that 30-40 % of confluency was obtained on the day of treatment with test compounds. On the day of compound treatment, cells were treated with the compounds or vehicle in fresh medium, and the plates were incubated for 72 hours in a 37 °C, 5% CO_2_. After 72 hours, the medium was discarded and CellTiter-Glo reagent was added to all wells (25 µl/well) according to the manufacturer (Promega, Cat # G7571). Plates were then incubated at 25°C in a thermomixer with shaking at 300 rpm. The luminescence signal was subsequently measured in a plate reader (Spectramax M5, Molecular Devices). Activity of the compounds was expressed as % inhibition, normalized between positive and negative controls. DRCs were fitted using a four-parameter fit equation.

### Mining and removal of known antimalarial scaffolds

To ensure chemical novelty, sub-structure searches in our internal database were conducted on the active hits to flag any scaffolds that are or have been in clinical development for use as an antimalarial drug. Fragments used in these searches are depicted in Table S3. The identification of biologically active compounds containing known antimalarial fragments was expected from the HTS. The presence of such compounds indirectly validates the HTS methodology. However, to increase the probability of identifying novel chemotypes and potentially novel malaria drug targets biologically active compounds containing known antimalarial fragments were deprioritized.

## Results

### Generation of a transgenic reporter parasite

In order to establish a sensitive and homogeneous assay for monitoring parasite replication in miniaturised format, we established a reporter parasite expressing a NanoLuc gene driven by the *ef1α* promoter (Figure 3A). The gene cassette was successfully introduced into *P. falciparum* NF54 parasites by transfection and integration in the 3’ region flanking the *pfs47* gene by single cross-over. The selected clone NF54-nanoGlo showed replication rates comparable to those of NF54 wildtype parasites whereas Dd2 parasites, used for orthogonal confirmation assays, showed slightly lower growth rates (Figure 3B). When NF54-nanoGlo parasites were seeded in 1536-well plates, luminescence intensity increased following 72 hours of incubation, depending on the starting parasitemia and hematocrit (Figure 3C). Based on these results, 1.2% parasitemia and 1.5% hematocrit for inoculating replication assays were used for the subsequent experiments. Under these conditions, parasite replication could be inhibited by compounds used in marketed antimalarial drugs (Figure 3D) although IC_50_ values of lumefantrine and piperaquine were slightly higher than previously reported ^17^. Parasites were not sensitive to antifolates such as pyrimethamine as a result of the integration of the *dhfr* selection cassette into the parasite’s genome (Figure 3D). Testing of a larger set of 123 antimalarial reference compounds in the nanoluciferase format and a previously established replication assay based on monitoring pLDH activity using a fluorescent substrate ^18^ showed an excellent correlation (R^2^ = 0.83) between the two assay formats.

With a view on establishing speed of kill assays, parasite replication by proxy of the increase in luminescence was monitored as a function of time for parasites treated with 10 µM dihydroartemisinin (DHA) or vehicle control. The results indicated a robust difference in signal (S/B > 2.0, Z’ > 0.5) with incubation times as short as 12 hours (Figure 3E).

### Test cascade overview

A comprehensive primary screening and hit triage assay cascade was designed and then used to identify and prioritize novel and chemically diverse high-quality hits from the MMV Hit Generation Library 1 (HGL1) (Figure 2). The library of 141,786 compounds was first tested in the primary screen at a single concentration (2 µM) against *P. falciparum* NF54 using a nanoGlo read-out. The activity of 4,596 compounds that passed the cut-off value of a Z-score lower or equal to -4 in primary screen was retested in the primary (*P. falciparum* NF54, nanoGlo) and an orthogonal assay (*P. falciparum* NF54, pLDH). 483 compounds that complied with the criteria of above 30% growth inhibition in at least one of the two assays were selected for further profiling. The selected compounds were tested in dose-response curve (DRC) assays for drug sensitive (*P. falciparum* NF54, nanoGlo and *P. falciparum* NF54 pLDH) and drug resistant parasite strains (*P. falciparum* Dd2, pLDH) and for cell viability (HepG2 cells). Parasite growth inhibition was measured after an incubation of 72 hours. Furthermore, the compounds were tested in a *P. falciparum* NF54 growth inhibition assay with a 12-hour incubation time. This assay was designed to evaluate the speed of action of the compounds and highlight compounds with potential to have a fast rate of parasite kill. Out of 75 compounds that fulfilled all of the ideal criteria (described in Figure 1), 46 were deemed as attractive in terms of chemical novelty relative to known antimalarial drugs and diversity based on Tanimoto similarity in 2D fingerprints. This set of compounds was then further complemented with additional compounds that complied with at least three of the five selection criteria that were set for DRCs. 76 compounds meeting the MMV Screening Active criteria, of which only 61 were commercially available in solid format for purchasing and retesting. To further validate the initial results, the efficacy of these compounds was retested against *P. falciparum* 3D7 in a pLDH assay and human HepG2 cells in a different laboratory, resulting in 33 confirmed active compounds with in IC_50_ lower than 2 µM in a whole cell asexual blood stage growth inhibition assay and higher than 10 µM in a cell viability assay from two independent test centers. Although direct activity comparison could not be performed since different strains, media and readouts were used, independent retention of activity greatly strengthened confidence in the quality of the hits and the robustness of the supporting data package.

**Figure 2:**
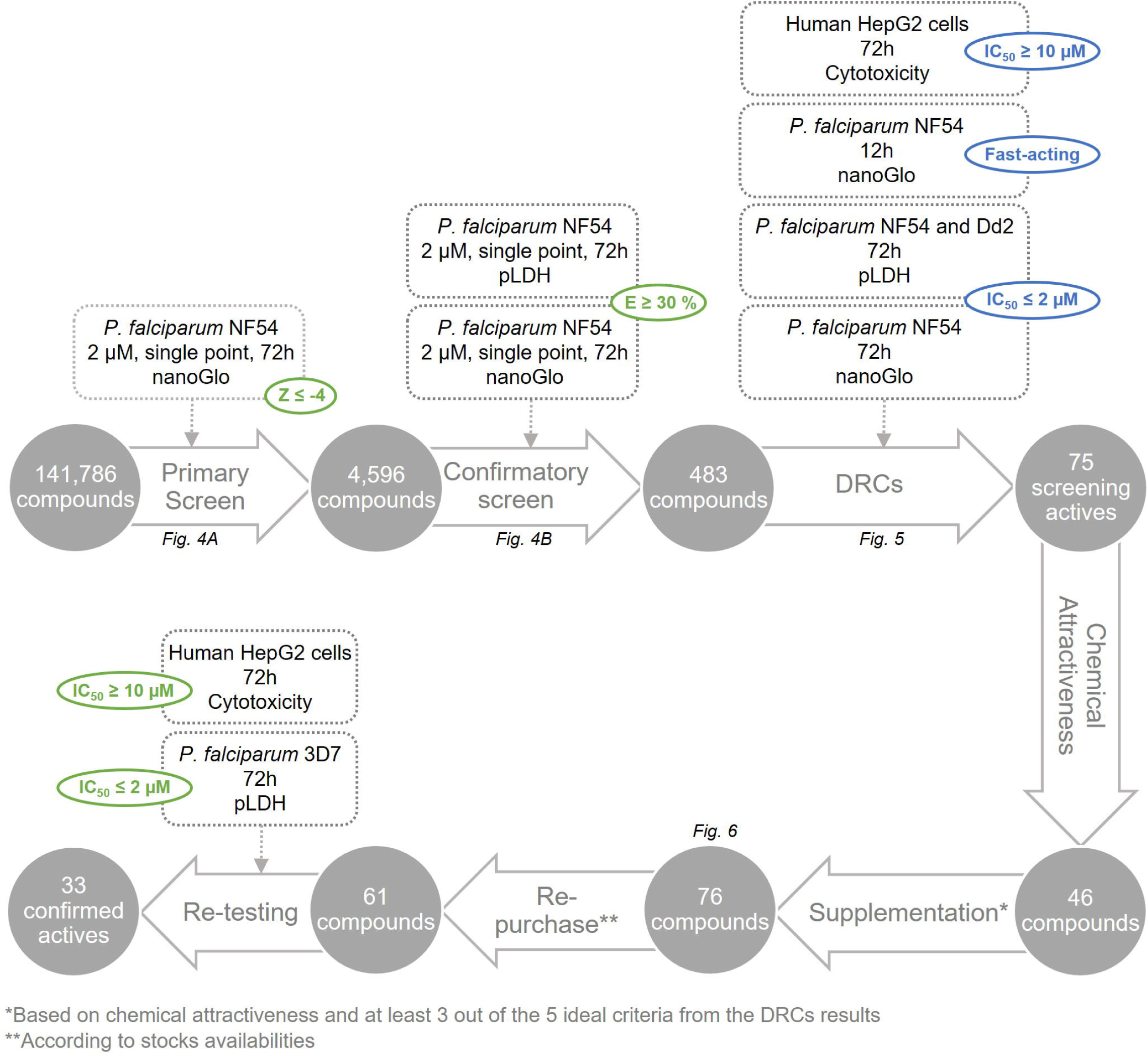
Generation and prioritization of actives from the Hit Generation Library 1 (HGL1). Selection cascade used to triage hits from HGL1. A primary screen was conducted on the full library, with a single concentration against *P. falciparum* NF54 using a nanoGlo read-out. 4,596 compounds passed the cut-off value of a Z-score lower or equal to -4 (highlighted in green) and were further retested in two orthologue assays (nanoGlo and pLHD read-outs). 483 compounds complied with the effect criteria of a minimum of 30% of growth inhibition activity (highlighted in green) in at least one of the two assays. These compounds were then subjected to five dose-responses assays (DRCs), each linked to ideal, although not absolutely required, criteria (highlighted in blue) : three parasitological assays (*P. falciparum* NF54 in both nanoGlo and pLDH read-outs and *P. falciparum* Dd2 in pLDH read-out using an incubation time of 72h), one assay crudely evaluating the speed of action of the compounds (*P. falciparum* NF54 in nanoGlo read-out using an incubation time of 12h), and a cytotoxicity assay against human HepG2 cells over 72h of incubation. Out of 75 screening actives fulfilling all of the ideal criteria, 46 only were deemed attractive in terms of chemical novelty and diversity. This set of compounds was then further complemented with others meeting at least three of the five DRCs criteria, as well as possessing chemical novelty and diversity, leading 76 selected compounds. Based on stock availabilities, 61 of these were re-purchased for retesting the efficacy against *P. falciparum 3D7* in a pLDH assay and human HepG2 cells at a different industry site. 33 compounds fulfilled the necessary criteria of an IC_50_ lower than 2 µM in 3D7 and higher than 10 µM in HepG2 and were qualified as Confirmed Actives. Strict selection criteria are depicted in green, and ideal ones on blue.

**Figure 3:**
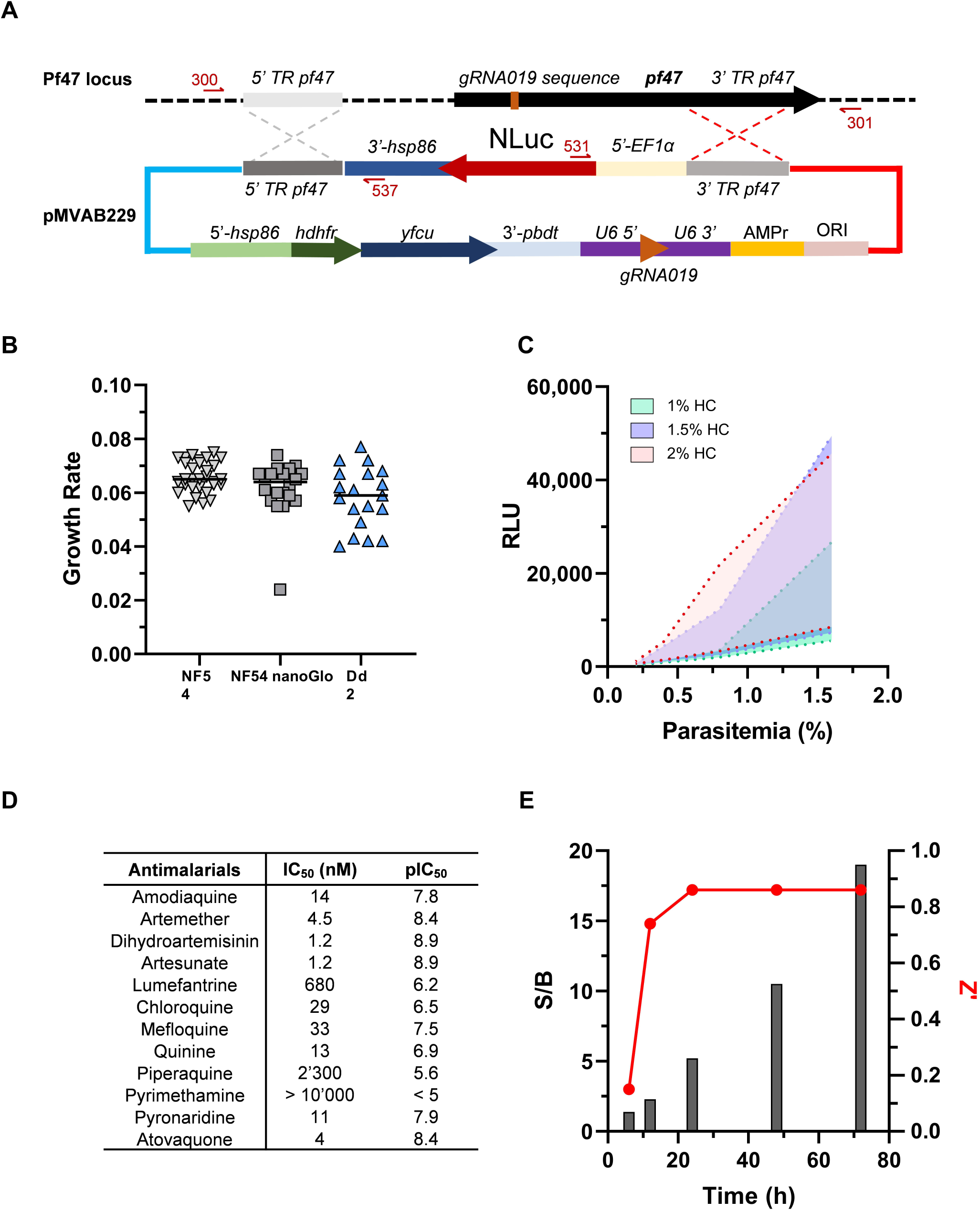
*P. falciparum* NF54 nanoGlo strain design, characterization and early validation for asexual blood stages High Throughput Screening. (A) Genomic targeting strategy for integration of a nanoLuc (Nluc) reporter gene driven by a *P. falciparum EF1a* promoter in the *Pf47* locus. The transgenic line used in these studies originated from a single cross-over, indicated by the red dashed cross. 5’-3’-TR: 5’ or 3’ end translated region; *gRNA019*: guide RNA sequence (not used here); *hsp86*: heat shock protein 86; *hdfhr*: human dihydrofolate reductase gene; *yfcu*: yeast cytosine deaminase/uridyl phosphoribosyl transferase gene, *3’pbdbt*: *P. berghei* terminator sequence.; AMPRr: Ampicilin resistance gene; ORI: origin of replication. (B) Growth rate of the *P. falciparum* NF54 nanoGlo strain compared to the NF54 and Dd2 strain, described as N(t) = N(0) e^(rt), N(t) the number of parasites at time t, N(0) starting parasitemia and r the growth rate. The median of individual measurements is displayed. (C) Initial optimization of parasitemia and hematocrit in 1536 well format growth assay. Mean of 32 observed luminescence signals in parasites treated with vehicle (0.1% DSMO, maximum) or 10 µM of DHA (minimum). Parasitemia and hematocrit (HC) were varied as indicated. (D) Response of the NF54 nanoGlo strain against marketed antimalarials. Mean of at least 2 independent replicates (E) Performance of the strain in 1536 well format growth assay as a function of time. Representative S/B and Z’-score values of one experiment are reported based on luminescence measurements of parasites treated with 10 µM of DHA or vehicle (0.1% DSMO) for various amount of time.

### Validation of the nanoGlo assay in 1536 wells format

Variation of incubation times (Figure S3A) showed that the signal to background of the nanoGlo assay was significantly higher with 3 days compared to the two days incubation (± 30 vs. 17). However, long incubation times resulted in clear plate edge effects due to evaporation. Therefore, the Z’ value of the plates including the edge wells were lower compared to Z’ values in absence of edge wells (0.5 vs 0.8). Both readout technologies (nanoGlo and pLDH) showed the similar plate patterns. Therefore, to obtain robust data across all plates, we used a 3-day incubation time and excluded the outter wells for HTS. This criteria led to the possibility of screening 1176 compounds per plate. The results of the incubation time (Figure S3B) clearly showed an increase in S/B and Z’ for the pLDH assay when incubated longer, with an optimum reached after approximately 30-45 minutes. The nanoGlo assay showed a steady decline in S/B and Z’ in time (Figure S3C), with an optimal incubation time shorter than 20 minutes. Both assays showed suffcient tolerance to DMSO with almost no effect up to 1% of DMSO final assay concentration (f.a.c.) (Figure S3D and F), above which resulted in decreased maximum signal and reduced S/B ratio. Overall, the nanoGlo assay showed robust results (Figure S3C and E). After optimizing the assay parameters, the robustness set compound collection ^16^ was tested using the optimal screening conditions. The robustness set consisted of a full 1536-well plate and a total of 262 compounds spread across various classes of compounds with assay interfering properties, clean compounds for which no obvious assay interfering properties were expected and DMSO controls (Figure S4A). For the analysis, the outer wells were removed from the dataset. Selecting active compounds based on a 30% effect cutoff or a Z-scores ≤-4 (Figure S4B) showed that the primary screening assay is relatively sensitive to aggregators, chelators, coloured compounds and redox modifiers, indicating the necessity for an orthogonal confirmation assay. The testing of the robustness set was combined with the testing of a set of known antimalarials. A total of 63 compounds were tested in 10 dose point DRCs using two different malaria strains (NF175 and NF54). The NF175 strain was tested using pLDH as readout and the NF54 strain was tested using nanoGlo as readout. The correlation between the two assays was excellent (R^2^ value of a linear regression of 0.978 for the 63 compounds) (Figure S5 and Table S4).

### Primary and confirmatory screens

A total of 141,786 compounds were screened using the optimized and miniaturized nanoGlo assay in two runs. Results of the screen were calculated using Activity Base XE and Dotmatics software. The S/B and Z’ values of all plates were in the acceptable range (from 11.6 to 23.1 and from 0.63 to 0.85, respectively, data not shown), and all plates passed the quality control criteria. The Z-score frequency distribution of all compounds is shown in Figure 4A. 4,596 compounds with a Z-score ≤-4 were considered stastistically active and were retested in both the nanoGlo and pLDH assays for activity confirmation. The inhibitory effects of these compounds was correlated (R^2^ value of a linar regression of 0.81) in both read-outs (Figure 4B). 483 compounds with at least 30% effect in one of the assays were selected for further profiling.

**Figure 4:**
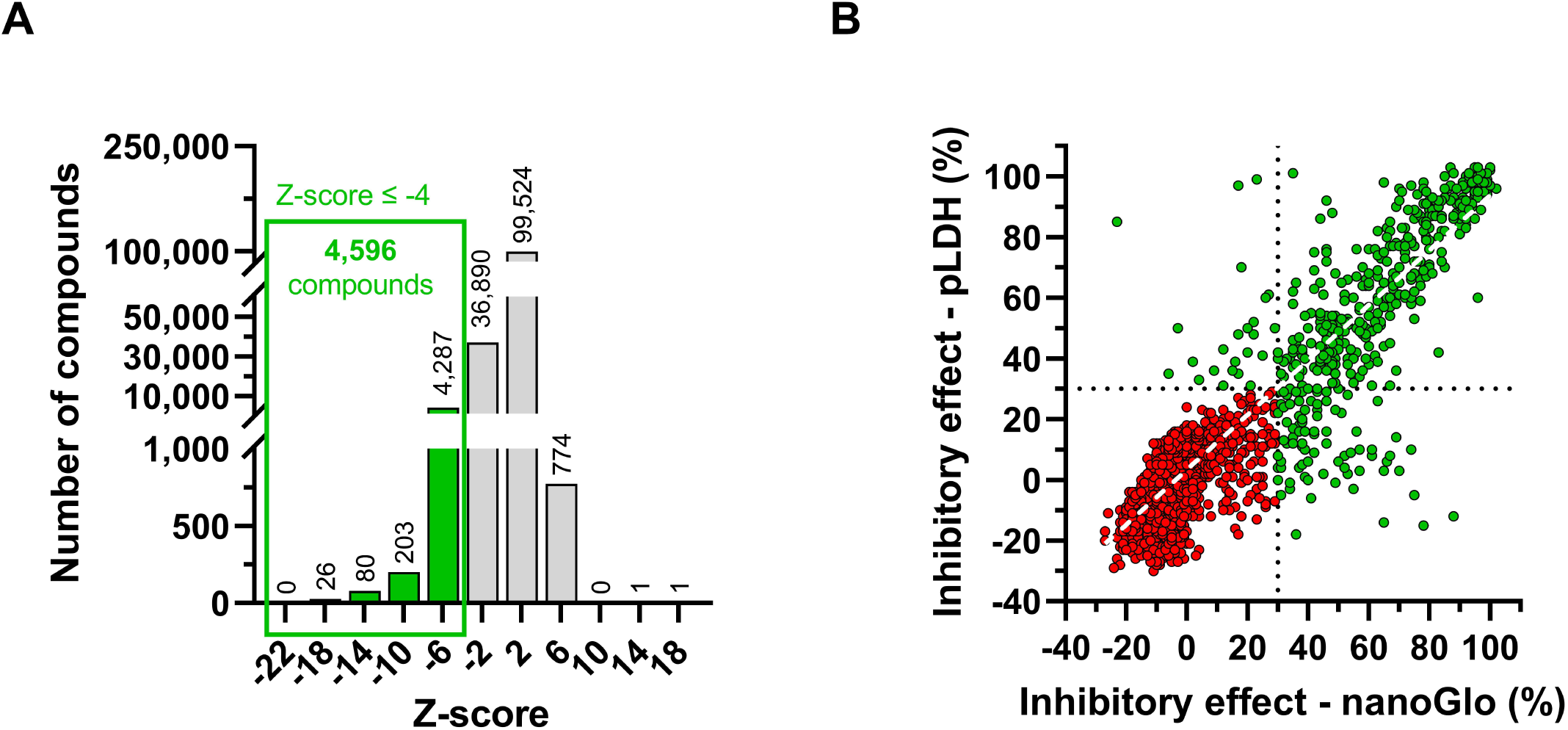
Primary and confirmatory screens results. (A) Z-score frequency distribution of the 141,786 of the Hit Generation Library in the primary nanoGlo screen. 4,596 compounds displayed a Z-score ≤ 4 (green) and were selected for activity confirmation. (B) Correlation between effects (%) in the pLDH and nanoGlo read-outs in compounds selected for activity confirmation. Of those, 483 compounds showing ≥ 30% inhibition effect in at least one of the two assays (in green) were selected for further dose-response characterization. A simple linear fit (white dashed line) between the two read-outs resulted in an R^2^ of 0.8124. The dotted lines represent the cut-off value of 30% effect.

### Screening actives

The 483 selected compounds were tested in multiple DRC assays to comprehensively characterize their pharmacology. Five different assays were carried out, including three parasite growth assays over 72 hours of incubation (*P. falciparum* NF54 in both nanoGlo and pLDH read-outs and *P. falciparum* Dd2 in pLDH read-out), one parasite growth assay over 12 hours of incubation (*P. falciparum* NF54 in nanoGlo read-out) in order to evaluate the speed of action, and a cytotoxicity assay against human HepG2 cells over 72 hours of incubation. Assays against NF54 in both read-outs reasonably correlated (R^2^ value of a linar regression of 0.6406, Figure 5B). As Dd2 is a multi-drug resistance strain compared to NF54 ^19^, the correlations between the outcomes of the assays against Dd2 and NF54 (nanoGlo and pLDH read-outs) were expectedly low (R^2^ values of a linar regressions of 0.1169 and 0.1662 respectively, Figure 5C and D). However, many compounds induced a response outside of the assay senstivity against Dd2, showing a markedly decreased sensitivity compared to NF54, and removing these points led to safisfactory correlations (R^2^ value of a linar regressions of 0.6406 in the nanoGlo read-out and 0.6030 in the pLDH read-out, Figure 5C and D). In line with previously described methods ^20^, slow- and fast-acting compounds were discriminated by comparing pIC_50_ values from 12 hour versus 72-hour incubation. To this end, we determined the slope of the change in pIC_50_ over time and used logistic regression to compare the slopes to Parasite Reduction Rate ^21^ data for a reference set of 20 compounds with known speed of action. The results show that the change in pIC_50_ discriminates well between fast and slow acting compounds (Figure S6). At a cut-off of ≤0.009587 for the slope of the change in pIC_50_ over time (i.e., less than a 4.9-fold increase in IC_50_ from 12 to 72 hours), 92% of the fast-acting compounds were correctly identified whereas the false positive rate (slow-acting compounds wrongly classified as fast) of 14% (Figure S6). Ideally, selected compounds should display IC_50_ values lower than 2 µM against NF54 (pLDH and Nano-Glo read-outs) and Dd2 and 72 hours of incubation, be fast-acting and have an IC_50_ value higher than 10 µM in human HepG2 cells (Figure 5A, left panel). Only 75 of the tested compounds met all these criteria, and considered screening actives, resulting into a hit rate of 0.05%. The number of compounds passing each individual criterion is displayed in Figure 5A (right panel).

**Figure 5:**
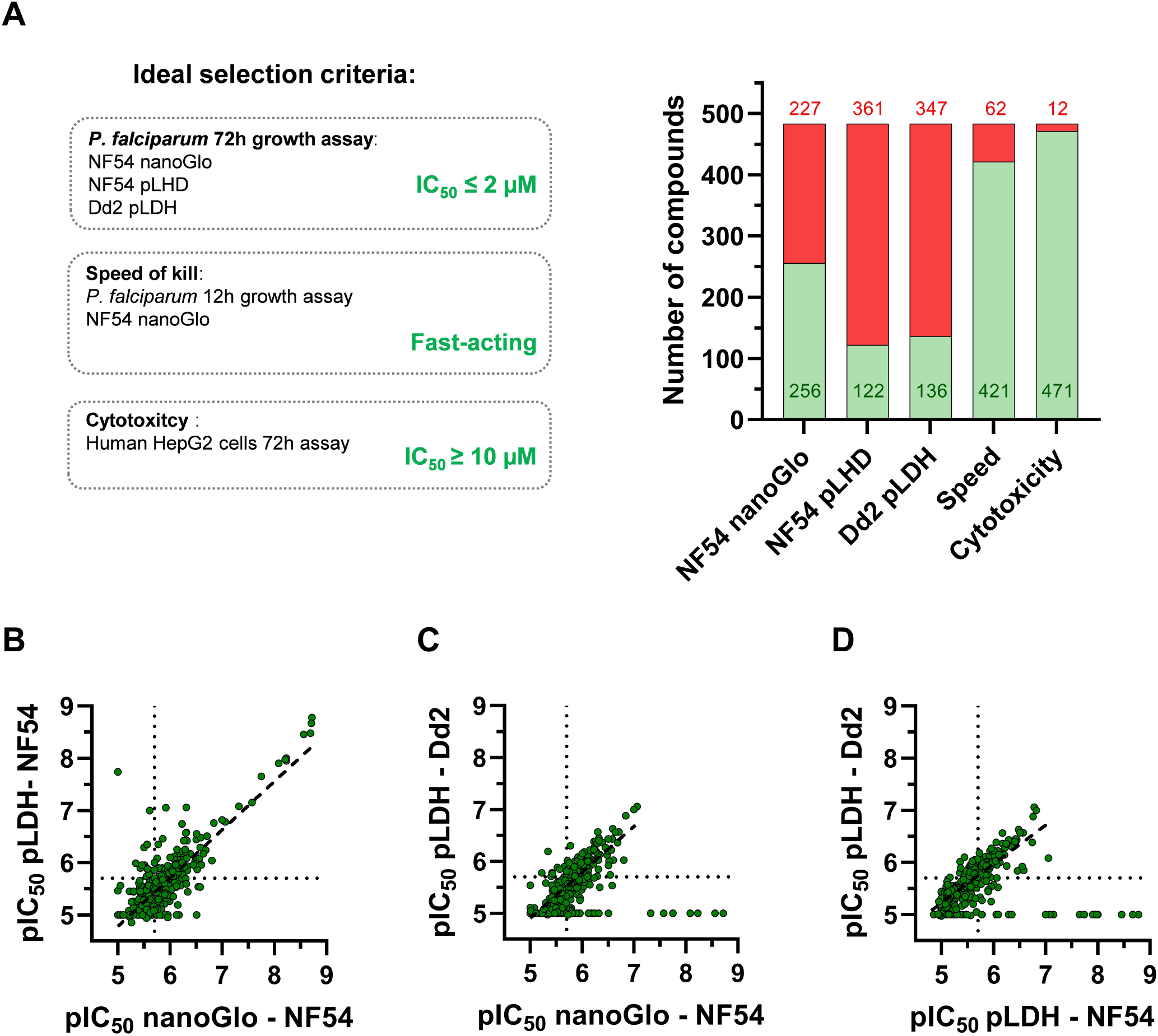
Outcome of the five DRC assays on the 483 selected compounds. (A) Left: ideal selection criteria: the IC_50_ values of the three classical parasitological assays (*P. falciparum* NF54 in both nanoGlo and pLDH read-outs and *P. falciparum* Dd2 in pLDH read-out using an incubation time of 72h) should reach a maximum of 2 µM; compounds should be fast acting based on a 12h and 72 assays comparison (See Fig. S5); a toxicity on human HepG2 cells up to 10 µM is accepted. Right: Number of compounds reaching (green) or failing (red) each criterion. (B) Correlation between the pIC_50_ values of each compound in the two read-outs at 72 h incubation (x-axis: nanoGlo; y-axis: pLDH). A simple linear fit (dashed line) between the two read-outs resulted in an R^2^ of 0.6406. (C) Correlation between the pIC_50_ values of each compound against the two *P. falciparum* strains at 72h incubation (x-axis: NF54, nanGlo; y-axis: Dd2, pLDH). A simple linear fit between the strains, including all the compounds would result in an R^2^ of 0.1169. However, by removing compounds with activity below the level of sensitivity of the assay (IC50 ≥ 10 µM), a simple linear fit (dashed line) between the two resulted in an R^2^ of 0.6030. (D) Correlation between the pIC_50_ values of each compound against the two *P. falciparum* strains (x-axis: NF54, pLDH; y-axis: Dd2, pLDH). A simple linear fit between the strains, including all the compounds would result in an R^2^ of 0.1662. However, by removing compounds with activity below the level of sensitivity of the assay (IC50 ≥ 10 µM), a simple linear fit (dashed line) between the two resulted in an R^2^ of 0.6840. The dotted lines represent the cut-off values of an IC_50_ value at 2 µM.

### Confirmed actives

Out of the 75 compounds that fulfilled the five criteria mentioned above, 46 were considered most compelling in terms of chemical diversity and chemical novelty with respect to known antimalarial chemotypes. They were then further complemented with additional compounds from the primary screen that complied with at least three of the five criteria, showed a NF54 vs. Dd2 IC_50_ ratio no higher than 3, and were structurally novel, leading to a set of 76 compounds. Although these compounds did not necessarily display an IC_50_ value lower than 2 µM against *P. falciparum* in the three assays (Figure 6A), all of them complied with the ideal value against *P. falciparum* Dd2 (Figure 6B). 72 compounds were considered fast acting and only 3 displayed an IC_50_ against HepG2 cells higher than 10 µM (Figure 6C). Out of these 76 compounds, 61 could be repurchased and new batches were retested against *P. falciparum* 3D7 (pLDH read-out, 72 hours of incubation) and HepG2 (72 hours of incubation) in a different testing center (Figure S7). Aggregation of the combined results allowed the identification of 33 compounds that met the MMV Confirmed Active criteria (https://www.mmv.org/research-development/information-scientists) as determined in 2 independent test centres (Table S5).

**Figure 6:**
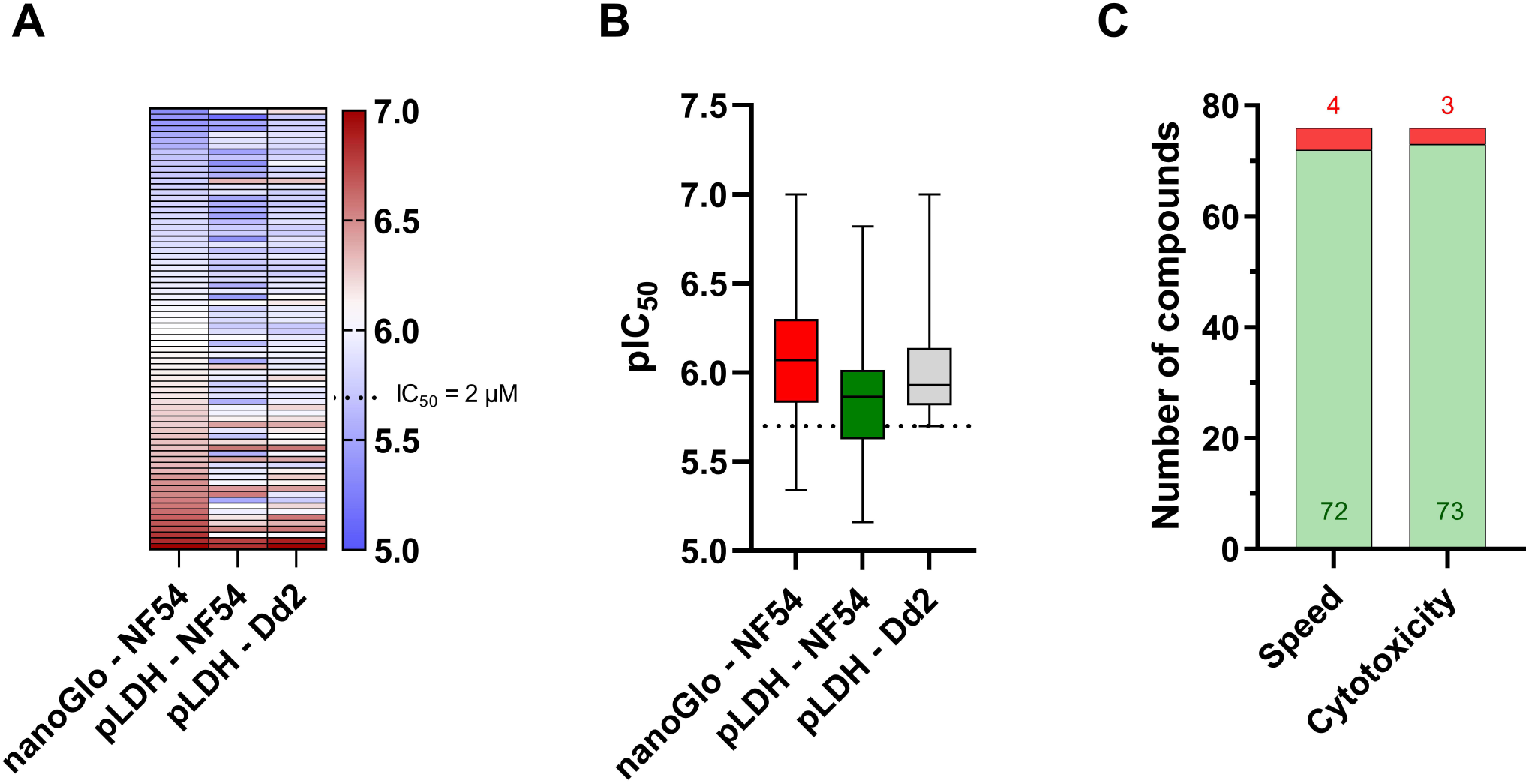
Characteristics of the 76 selected compounds, based on the DRCs results. (A and B) pIC_50_ values of each screening active in the three classical parasitological assays using an incubation time of 72h: *P. falciparum* NF54 in both nanoGlo and pLDH read-outs and *P. falciparum* Dd2 in pLDH read-out, as individual values (A, heatmap) and as a whole (B, boxplot. Median, 25 and 75% percentile and minimum and maximum values represented). The dotted line represents the cut-off values of an IC_50_ value at 2 µM. (C) Number of compounds reaching (green) or failing (red) the speed of action and cytotoxicity criteria.

### Visualization of the chemical space and examples of Confirmed Actives

A t-distributed stochastic neighbour embedding (SNE) analysis was conducted to visualize the chemical space occupied by the confirmed actives. Figure 7 depicts the chemical space plot of the whole library and the confirmed actives with respect to the launch drug space. Known antimalarial drugs have been added to this plot as reference points. The proximity of two points represents the structural similarity between the corresponding compounds using a Tanimoto index based on a 2D path-based fingerprint. The distribution of points is defined by approximately 1,300 launched small molecule drugs using the t-distributed stochastic neighbour embedding algorithm ^22^. The chemical structures of 3 representative confirmed actives are also depicted in Figure 7: MMV1669145, MMV1747770 and MMV1722301; and their biological, physicochemical and ADMET properties are displayed in Table S6. Moderate to good aqueous solubility was observed (>13 µM).

**Figure 7:**
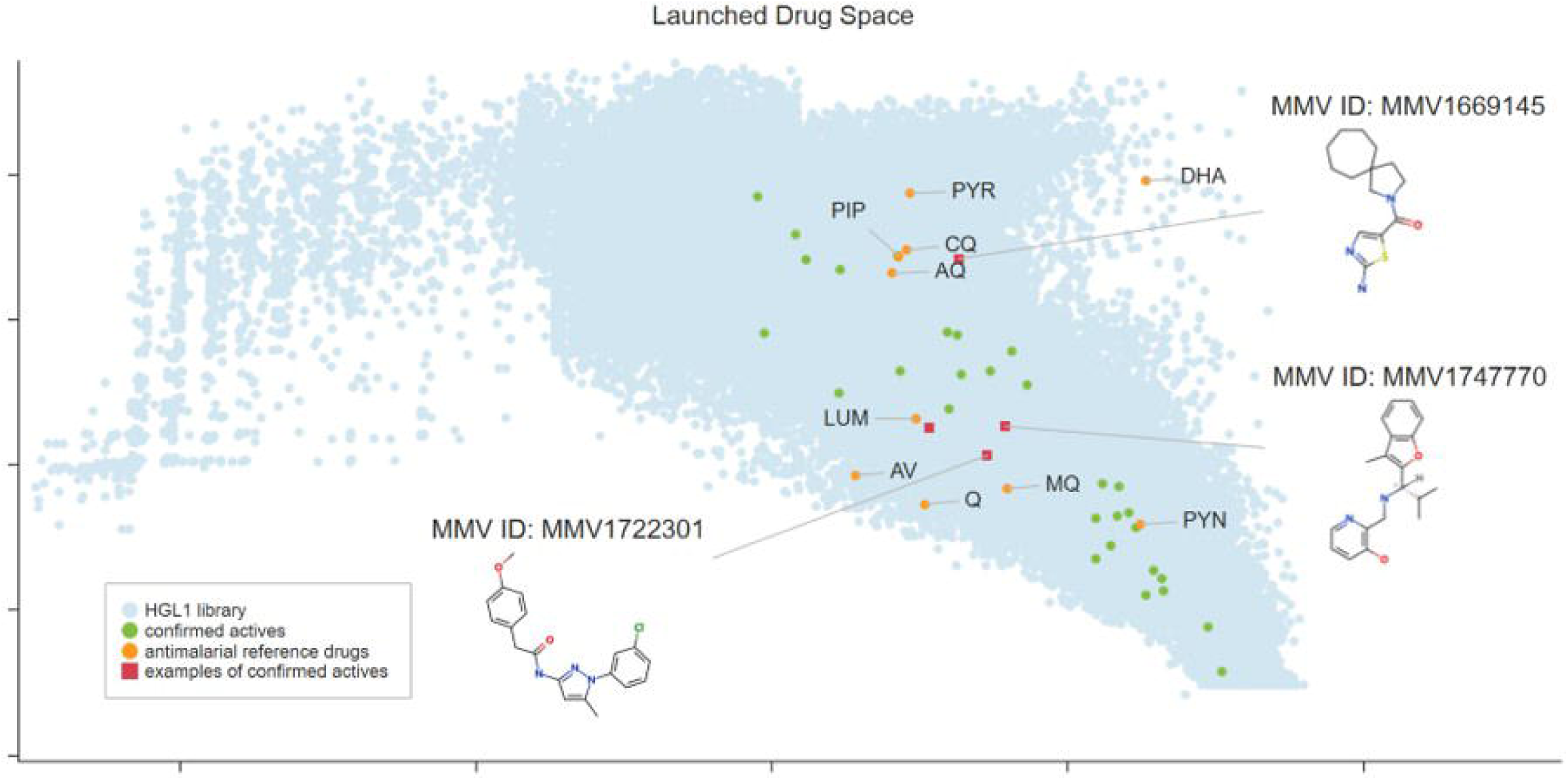
Summary of the drug space distribution. Illustration of the chemical space covered by the active compounds (green) and antimalarial reference compounds (orange) compared with the HGL1 library (light blue points). Here, the proximity of two points represents the structural similarity between the corresponding compounds defined using a Tanimoto index based on a 2D path-based fingerprint. The distribution of points is defined by approximately 1,300 launched small molecule drugs using the t-distributed stochastic neighbor embedding algorithm. DHA: dihydroartemisinin; PYR: pyrimethamine; PIP: piperaquine; CQ: chloroquine; AT: atovaquone; AQ: amodiaquine; LUM: lumefantrine; Q: quinine; MQ; mefloquine; PYN: pyronaridine.

## Discussion

Phenotype-based screens have shown great value in identifying first-in-class small antimalarial molecules in comparison to other screening methods ^23^ and the two last decades have seen promising antimalarial clinical candidates emerge from such large phenotypic screens ^24^. The GSK screen ^10^, in which approximately 2 million compounds were tested against *P. falciparum* blood stages with a hit rate around 0.56%, resulted in the current preclinical candidate MMV367, that will shortly enter clinical phase I. The SJCRH ^7^ and GNF ^9^ screens, with equivalent screening active hit rates of approximately 0.05% and 0.03% respectively, led to the identification of SJ733 ^25^ and KAE609 ^26^, both PfATP4 inhibitors, both currently progressing though clinical phases ^27^ (Clinical Trial ID: NCT04709692 and NCT04675931, respectively). Despite the novelty of the chemical scaffold and modes of action at the time, the risk of clinical resistance development of PfATP4 inhibitors is now widely recognized ^27^ highlighting the necessity of a clear focus on increased diversity in the next phenotypic screens. Therefore, in this study, care was taken during the library design to exclude compounds structurally related to compounds present in either the MMV in-house database or the ChEMBL-NTD archive.

While high-throughput screening of large diversity-based compound collections is a versatile approach to rapidly explore drug-like chemical spaces and identify bioactive molecules, a major challenge after primary screening and confirmation of activity is designing an appropriate follow up hit selection assay cascade to triage compounds with undesired modes of action and readout interfering properties. A scalable approach to identify such liabilities in primary screening assays is a chemical biology one in which HTS-ready assay is perturbed with various classes of known assay interfering properties ^28–30^ to make informed decisions for setting active selection criteria and employ appropriate follow up assays. Using the robustness set compound collection (Figure S4A), we identified liabilities mainly towards redox cycling compounds, coloured compounds, aggregators and chelators when activity selection criterion was set to z-score values below or equal to -4 (Figure S4B). Raising the bar and setting a cut-off of >30% effect resulted in a drastic reduction of liabilities towards redox cycling compounds, moderate reduction of liabilities towards coloured and metals and slightly less liabilities towards auto-fluorescent compounds (Figure S4B). To this end, for the primary screen we set z-score values below or equal to –4 as an active selection criterion to unbiasedly select all statistically significant compound activities resulting in 4,956 compounds. To minimize the risk of selecting compounds with undesired modes-of-action and/or readout interfering properties, on the one hand during active confirmation phase, we used a cut-off of above or equal to 30% effect in the primary assay and used an orthogonal pLDH assay and on the other, we included a HepG2 cytotoxicity assay during DRC testing and employed cheminformatic filtering of compounds with undesired properties.

In an effort to enhance sensitivity in parasitic phenotypic screening, we used a nanoluciferase reporter, engineered from a deep-sea shrimp luciferase enzyme and which provided an extremely bright bioluminescent signal with a long half-life ^31^. Given that an asynchronous parasite culture was used, in which at all times a fraction of the parasites is replicating, we could detect a robust difference in luminescence signals between drug-treated and untreated parasites with incubation times as little as 12 hours. Previous work using [^3^H]hypoxanthine incorporation assays has shown that compounds with rapid speed of action show a relatively minor shift in potency when comparing data from 24-hour incubations with those of 72-hour incubations, whereas slow-acting compounds show a larger difference ^20^. Here we have further simplified this analysis by providing a non-radioactive assay amenable to high throughput screening in 1536 well plates. In addition, we have further evaluated the change in IC_50_ over time in relation to killing rates. Our analyses of 20 compounds with known killing rates show that our luminescence-based IC_50_ determinations provide a straightforward classification of fast and slow killing compounds with excellent specificity and sensitivity. A fast mode of action is a desired property, as it provides rapid parasite clearance and relief of symptoms in malaria treatment, and a lower risk of the emergence and selection of resistance mutations.

Given that we used progressively stringent selection criteria, the screening active hit rate (0.05%) observed in this study is relatively low, although it still falls in the same range as the SJCRH and GNF screens. Strong emphasis was put on identifying compounds with both structural novelty and high quality, generating a comprehensive data package in order to identify attractive new starting points for antimalarial drug discovery projects. Therefore, stringent cut-off criteria were applied (five selection criteria for DRCs, IC_50_ < 2 µM) and diligent data mining was performed to remove known antimalarial scaffolds. However, to avoid discarding attractive chemotypes as a result of the strictness of these criteria, selected hits fulfilling at least three of the five selection criteria were rescued. Re-synthesis and/or repurchase of the screening actives allowed independent confirmation of the structure and associated biological activity of the compounds but also to assess their chemical tractability. Moreover, performing this re-test in a different testing centre using a different read-out provided further confidence in the reliability of the observed antimalarial potency and quality of the hits. Early ADMET and physicochemical data package (LogD, human microsomal stability, rat hepatocytes stability, aqueous solubility, hERG inhibition) were collected on all the confirmed actives in order to help prioritization of the confirmed actives for follow-up in drug discovery projects and to assess any liabilities that would need to be addressed as part of the hit-to-lead optimization process.

The identification of new high quality, drug-like, chemical starting points is dependent on the quality of the compounds tested and the screening cascade. Malaria drug discovery projects cannot afford the luxury of testing compounds with inherent developmental challenges based on either a lack of structural novelty or chemical groups associated with poor pharmacokinetics or safety. Furthermore, prioritization of the compounds with antimalarial activity for follow up is greatly aided by a robust and reproducible data package with early assessment of both drug resistance and speed of kill. With this in mind, we have developed exacting compound selection criteria that we share with the wider community for improving the diversity and property profile of screening libraries. In line with MMV’s commitment to openness, where possible, through MMVopen and public databases all screening and structural data discussed are freely available. This study is the first in a planned series of related publications describing our efforts to generate new chemical staring points for antimalarial drug discovery projects and target identification with potential benefit across the malaria, and more generally infectious diseases, drug discovery community.

## Data availability

Screening and validation data sets have been deposited into ChEMBL-NTD (doi:10.6019/CHEMBL4888484).

## Acknowledgments

We recognize and appreciate the helpful contribution of Ronald ten Berge (PPSC). We gratefully acknowledge the technical contribution of the TCGLS team for the confirmatory assays.

