## Supplements for "Replenishing the malaria drug discovery pipeline: Screening and hit evaluation of the MMV Hit Generation Library 1 (HGL1) against asexual blood stage *Plasmodium falciparum*, using a nano luciferase reporter read-out"

### Figure S1

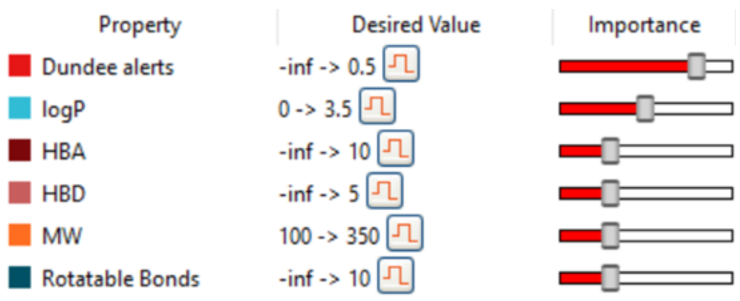


**Figure S1: Multi parameter optimization (MPO) score calculations, by Optibrium.** The molecular weight (MW), partition coefficient (logP), number of hydrogen bond donors (HBD) and acceptors (HBA) and number of rotatable bonds are calculated using StarDrop. Dundee alerts are provided by a model that accompanies the scoring profile [1]. The output is a score on a scale of 0 to 1 indicating the likelihood of achieving an ideal outcome for each property. More information on probabilistic scoring can be find in the StarDrop Reference Guide: https://www.optibrium.com/downloads/StarDrop_Reference_Guide_v6.3.pdf.

(1) Brenk, R.; Schipani, A.; James, D.; Krasowski, A.; Gilbert, I. H.; Frearson, J.; Wyatt, P. G. Lessons Learnt from Assembling Screening Libraries for Drug Discovery for Neglected Diseases. *ChemMedChem* **2008**, 3 (3), 435–444. https://doi.org/10.1002/CMDC.200700139.

#### Figure S2

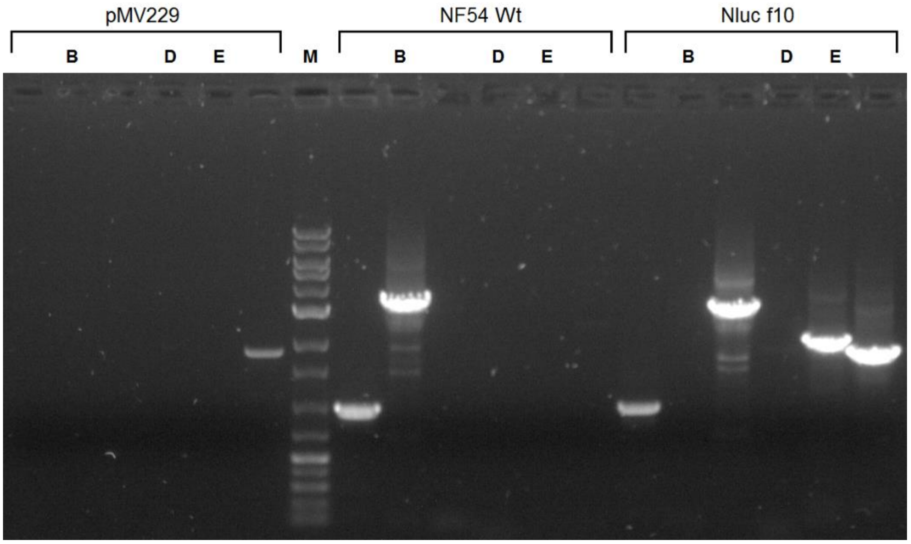


**Figure S2: Confirmation of integration of the Nanoluc reporter in the *P. falciparum* NF54 genome.** The figure shows agarose gel electrophoresis results for PCR products originating from the templates indicated on top of the figure. pMV229: plasmid used for transfection, NF54 Wt: untransfected NF54 wildtype parasites, Nlucf10: luciferase positive clone. PCR was performed using the following primer pairs: B) MWV 300/MWV 301 resulting in a 3.3 kb fragment for Wt, D) MWV 300/MWV 537, resulting in a 1.9 kb fragment in case of integration at the 5' end of the Pf47 gene, E) MWV 531/MWV 301, resulting in a 2.1 kb fragment in case of integration at the 3' end of the Pf47 gene. Lanes without a letter annotation show unrelated PCR results not relevant to this paper. Lane M shows a DNA size marker with bright bands at 3.0 and 0.5 kb.

#### Figure S3

A

| | pLDH | | nanoGlo | |
|---|---|---|---|---|
| | S/B | Z'value | S/B | Z'value |
| 2 days | 4.57 | 0.47 | 17.78 | 0.59 |
| 2 days | 3.75 | 0.18 | 15.59 | 0.32 |
| 2 days without edge | 4.62 | 0.63 | 18.15 | 0.64 |
| 2 days without edge | 3.78 | 0.31 | 15.71 | 0.31 |
| 3 days | 6.01 | 0.32 | 42.53 | 0.38 |
| 3 days | 5.91 | 0.33 | 25.69 | 0.57 |
| 3 days without edge | 6.24 | 0.73 | 44.81 | 0.74 |
| 3 days without edge | 6.13 | 0.79 | 26.37 | 0.83 |

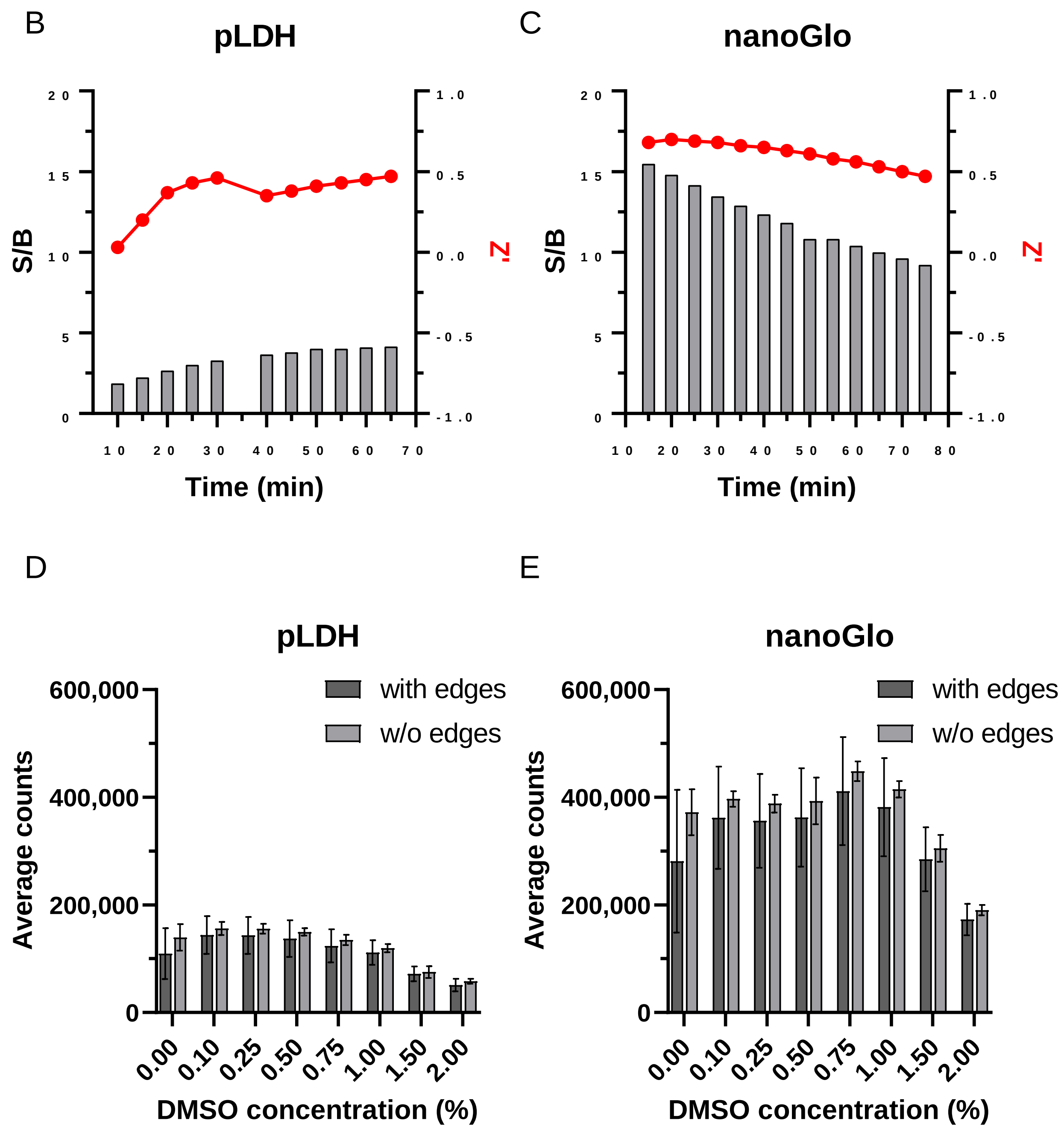

#### Figure S3

**Figure S3: Feasibility study performed at Pivotpark Screening Center.** (A) Performance of the pLDH and nanoGlo assays during a two- or three-days feasibility study, with or without including the edges of the 1526-well plates. (B and C) Stability of the pLDH (B) and nanoGlo (C) reagents in the respective assays over two days of incubation. The time in minutes after addition (X-axis) was plotted against two assay parameters: signal to background ratio (S/B) (left Y-axis, grey bar graph) and Z'-score (right Y-axis, red dots). (D and E) DMSO tolerance of the pLHD (D) and nanoGlo (E) assays. The final concentration (f.a.c.) of DMSO (%) was plotted against the average counts. DMSO tolerance was determined including (dark grey) or excluding (light grey) edge wells of the 1536-well plates, using DHA (0.8 mM) and other references compounds (1 and 10 mM).

All the assays were performed using dispensing equipment (Multidrop combi, ECHO and Certus) with 0.25% DMSO, 2 µM DHA, 1.2% parasitaemia, 1.5% RBC and 1:4 Nano-Glo (f.a.c) or pLDH (f.a.c.) and measured with a gain of 147 (pLDH) or 3600 (nanoGlo) and a focal height of 8.5 mm (pLDH) or 9.5 mm (nanoGlo). The plate was incubated two or three days in an incubator at 37°C and 5% $CO_2$.

**Figure S4**

A

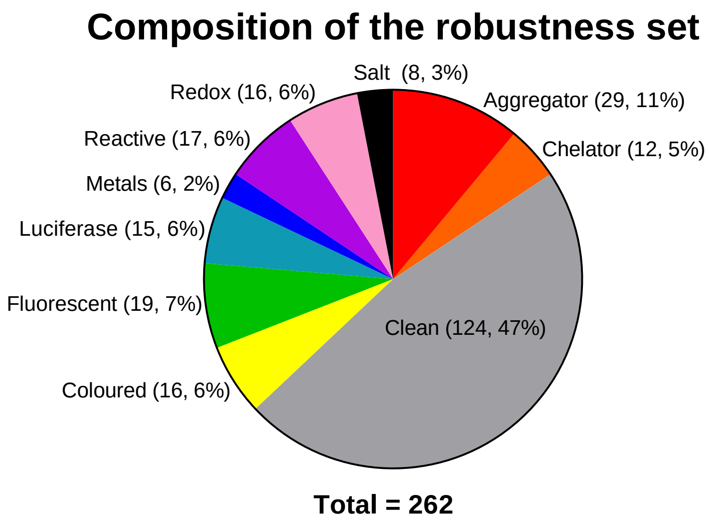


B

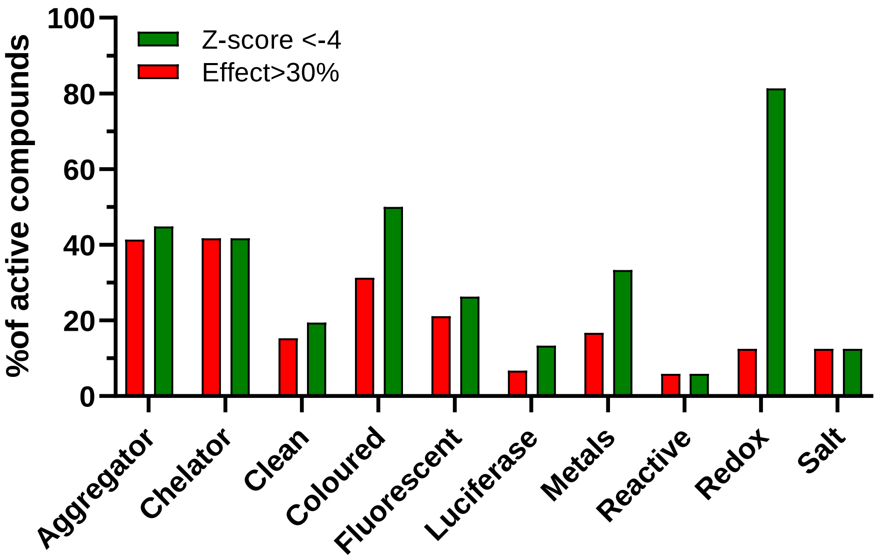


**Figure S4: NanoGlo assay robustness performed at Pivotpark Screening Center.** (A) Composition of the robustness set. (B) Robustness set at three-day incubation. Compounds were tested at 2 µM. The proportions of compounds above the effect and Z-score are represented. All the assays were performed in 1536-well plates using dispensing equipment (Multidrop combi, ECHO and Certus) with 0.25% DMSO, 2 µM DHA, 1.2% parasitaemia, 1.5% RBC and 1:4 Nano-Glo (f.a.c) and measured with a gain of 3600 and a focal height of 9.5 mm. The plate was incubated three days in an incubator at 37°C and 5% $CO_2$.

#### Figure S5

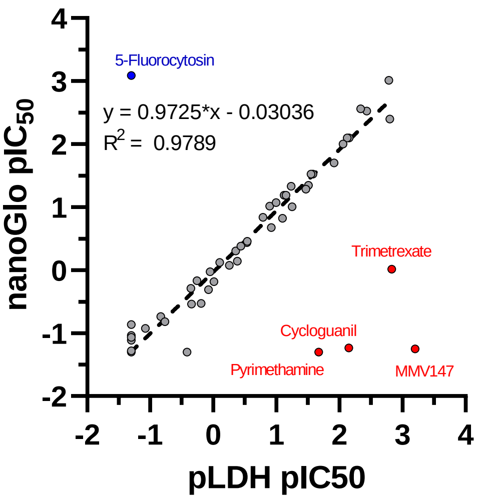


**Figure S5: Correlation between the two orthogonal assays performed at PPSC using the MMV Assay Validation box.** 63 compounds especially selected by MMV to validate the various assay platforms were tested in both *P. falciparum* NF175 with a pLDH read-out (x-axis) and *P. falciparum* NF54 nanoGlo (y-axis), in a 1536-well plate format. A few compounds particularly stood-up: four DHFR inhibitors (red) showed no activity in the NF54 nanoGlo strain, due to the presence of a DFHR resistance cassette as a remnant of the genome editing process.; and 5-Fluorocytosin, interfering with the pLDH fluorescent read-out, leading to a signal above the sensitivity of the assay. Despite these five outlier compounds, the correlation bwteeen the two assay was excellent, leading a $R^2$ value of a simple linear fit (dashed line) of 0.9789.

**Figure S6**

A

| Clasification based on: | | |
|---|---|---|
| | *PRR* | *Category* |
| MMV000130 | 0 | 1 |
| MMV018052 | 1 | 1 |
| MMV019721 | 1 | 1 |
| MMV1542805 | 1 | 1 |
| MMV1582617 | 1 | 1 |
| MMV390048 | 0 | 0 |
| MMV667546 | 0 | 0 |
| MMV673157 | 1 | 1 |
| MMV674849 | 1 | 1 |
| MMV674996 | 0 | 0 |
| MMV689134 | 0 | 0 |
| MMV892646 | 0 | 1 |
| MMV892941 | 0 | 0 |
| MMV979379 | 0 | 1 |
| Chloroquine | 1 | 1 |
| Piperaquine | 1 | 1 |
| Pyronaridine | 1 | 1 |
| MMV390048 | 0 | 0 |
| SJ733 | 0 | 1 |
| MMV669059 | 0 | 0 |

B

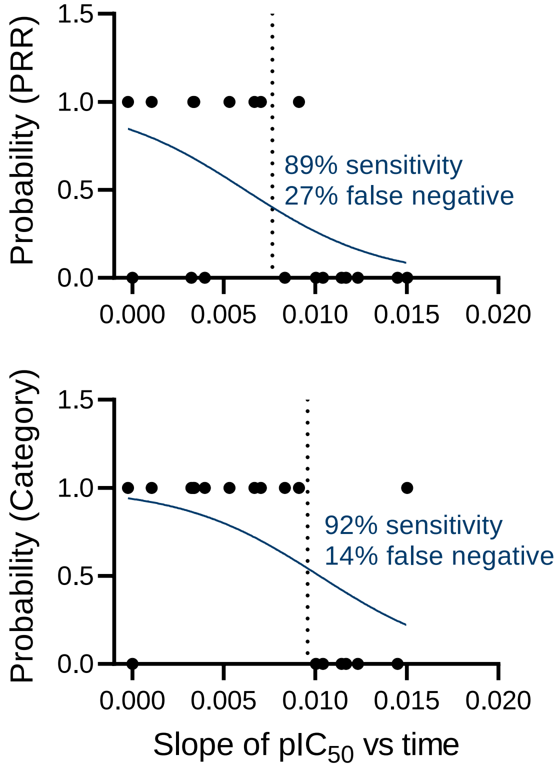


C

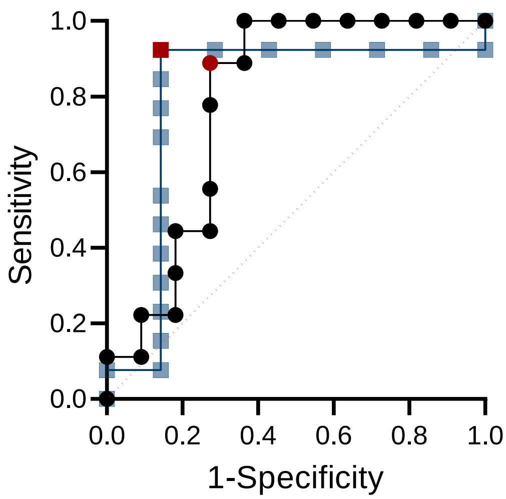


D

| Classification based on: | | |
|---|---|---|
| | *PRR* | *Category* |
| Sensitivity (%) | 89 | 92 |
| False positive rate (%) | 27 | 14 |
| AUC of ROC | 0.7879 | 0.8022 |
| $pIC_{50}$ slope cut-off | 0.007660 | 0.009587 |

### Figure S6

**Figure S6: Analysis of compounds potency as a function of incubation time to discriminate between slow- and fast- killing screened compounds.** (A) A set of reference compounds was categorized as fast(1)- or slow(0)-acting according to their PRR value (Fast = PRR >3.5; Slow = PRR ≤ 3.5) or their categorization by expert observation of their full PRR curve. (B) This classified compounds were plotted against the change in potency ($pIC_{50}$) over time, for both methods of classification. The solid line indicates the result of a simple logistic regression analysis. (C) Receiver Operating Characteristic (ROC) of the simple logistic regression analysis, showing the sensitivity (rate of true positive classification, i.e. correct classification of fast acting compounds) against 1-specificity (rate of false positive, i.e. the incorrect classification of slow-acting compounds as fast-acting) for both methods of classification (PRR value in black, categorization in blue). Selected points are highlighted in red. (D) Summary of the simple logistic regression and ROC parameters and cut-off value used to discriminate the screened compounds between slow – and fast-acting for both method of classification. For the purpose of the classification of the 483 confirmed hits, the cut-off value obtained with the expert categorization method was used.

#### Figure S7

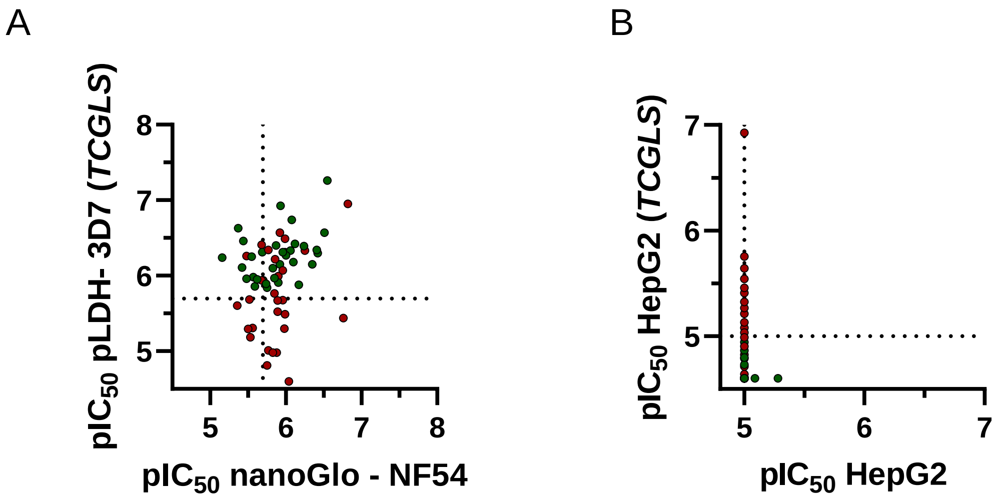


**Figure S7: Comparison of the 61 re-purchased compounds cross-tested at TCGLS in parasitological and toxicity assays.** (A) $pIC_{50}$ values of each compound against *P. falciparum* at 72h incubation using a pLDH read-out. (B) $pIC_{50}$ values of each compound against HepG2 cells. The 33 confirmed active compounds are represented in green.

**Table S1:** Hit Generation Library 1 characteristics.

| Molecular Weight | | LogP | | Alert | | MPO Score | |
|---|---|---|---|---|---|---|---|
| *Range (g/mol)* | *Number of compounds* | *Range* | *Number of compounds* | *Score* | *Number of compounds* | *Range* | *Number of compounds* |
| 0 - 225 | 3879 | (-)2.25 – (-)1.75 | 2 | 0 | 106596 | 0.025 – 0.075 | 28 |
| 225 – 275 | 20315 | (-)1.75 - (-)1.25 | 8 | 1 | 30185 | 0.075 – 0.125 | 67 |
| 275 – 325 | 39789 | (-)1.25 – (-)0.75 | 109 | 2 | 4190 | 0.125 – 0.175 | 144 |
| 325 – 375 | 43122 | (-)0.75 – (-)0.25 | 434 | 3 | 1176 | 0.175 – 0.225 | 554 |
| 375 – 425 | 20088 | (-)0.25 – 0.25 | 1748 | 4 | 410 | 0.225 – 0.275 | 315 |
| 425 – 475 | 9464 | 0.25 – 0.75 | 4953 | 5 | 89 | 0.275 – 0.325 | 673 |
| 475 – 525 | 4229 | 0.75 – 1.25 | 10528 | 6 | 41 | 0.325 – 0.375 | 1425 |
| 525 – 575 | 1244 | 1.25 – 1.75 | 17825 | 7 | 14 | 0.375 – 0.425 | 2446 |
| 575 – 625 | 476 | 1.75 – 2.25 | 24622 | 8 | 9 | 0.425 – 0.475 | 1791 |
| 625 – 675 | 107 | 2.25 – 2.75 | 26624 | 9 | 31 | 0.475 – 0.525 | 2393 |
| 675 – 725 | 22 | 2.75 – 3.25 | 22661 | 10 | 8 | 0.525 – 0.575 | 2890 |
| 725 – 775 | 10 | 3.25 – 3.75 | 14975 | 11 | 4 | 0.575 – 0.625 | 5101 |
| 775 – 825 | 6 | 3.75 – 4.25 | 8689 | 12 | 2 | 0.625 – 0.675 | 3339 |
| 825 – 875 | 2 | 4.25 – 4.75 | 4655 | | | 0.675 – 0.725 | 5285 |
| 875 – 925 | 1 | 4.75 – 5.25 | 2390 | | | 0.725 – 0.775 | 7476 |
| 925 – 975 | 0 | 5.25 – 5.75 | 1313 | | | 0.775 – 0.825 | 21534 |
| 975 – 1025 | 0 | 5.75 – 6.25 | 665 | | | 0.825 – 0.875 | 5740 |
| 1025 – 1075 | 0 | 6.25 – 6.75 | 320 | | | 0.875 – 0.925 | 8726 |
| 1075 – 1125 | 0 | 6.75 – 7.25 | 126 | | | 0.925 – 0.975 | 15461 |
| 1125 – 1175 | 0 | 7.25 – 7.75 | 71 | | | 0.075 – 1.00 | 57367 |
| 1175 – 1225 | 0 | 7.75 – 8.25 | 20 | | | | |
| 1225 - 1275 | 1 | 8.25 – 8.75 | 10 | | | | |
| | | 8.75 – 9.25 | 3 | | | | |
| | | 9.25 – 9.75 | 3 | | | | |
| | | 9.75 – 10.25 | 1 | | | | |

| Chemical bonds characteristics | | | | | |
|---|---|---|---|---|---|
| *HBD* | *Number of compounds* | *HBA* | *Number of compounds* | *Rotable* | *Number of compounds* |
| 0 | 40928 | 1 | 230 | 0 | 253 |
| 1 | 66109 | 2 | 2128 | 1 | 1496 |
| 2 | 30002 | 3 | 9201 | 2 | 6633 |
| 3 | 5308 | 4 | 23250 | 3 | 15176 |
| 4 | 391 | 5 | 35836 | 4 | 24068 |
| 5 | 15 | 6 | 33739 | 5 | 28718 |
| 6 | 2 | 7 | 21837 | 6 | 27043 |
| | | 8 | 11011 | 7 | 18557 |
| | | 9 | 3884 | 8 | 10772 |
| | | 10 | 1230 | 9 | 5085 |
| | | 11 | 315 | 10 | 2639 |
| | | 12 | 79 | 11 | 1248 |
| | | 13 | 14 | 12 | 625 |
| | | 14 | 1 | 13 | 246 |
| | | | | 14 | 111 |
| | | | | 15 | 44 |
| | | | | 16 | 26 |
| | | | | 17 | 7 |
| | | | | 18 | 5 |
| | | | | 19 | 3 |

**Table S2:** List of oligonucleotides used in construction of the reporter parasite strain.

| **Primer ID** | **Primer Sequence** |
|---|---|
| MWV300 | 5’-TACATTCAAATAACTCAGAGGGTAAC-3’ |
| MWV301 | 5’-GTTTGTGTATATTTACCTTACATTTATCTCC-3’ |
| MWV524 | 5’-AATTCTCGAGTCGACTTTTAACGGTTCACCCCTCTTAACC-3’ |
| MWV525 | 5’-ATCACCATGGAATAGTTTAGTATATTAATATATATG-3’ |
| MWV531 | 5’-CAGGACAATCCTTTGGATCG-3’ |
| MWV537 | 5’-TCATCAAATAAGGTAGCCGGC-3’ |
| MWV539 | 5’-TCGACAAGCTTGGGCCCGTACGCCGC-3’ |
| MWV540 | 5’- GGCGTACGGGCCCAAGCTTG-3’ |

**Table S3:** Examples of known antimalarial scaffolds

| Known antimalarial scaffold | Tag | Known antimalarial scaffold | Tag |
|---|---|---|---|
| 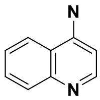 | 4-Aminoquinolines | 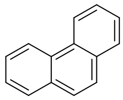 | Halofantrine |
| 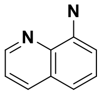 | 8-Aminoquinolines | 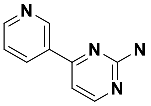 | Imatinib |
| 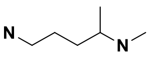 | Aminoquinolines side-chain | 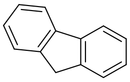 | Lumefantrine |
| 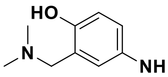 | Amodiaquine | 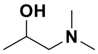 | Lumefantrine side-chain |
| 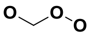 | Artemisinin | 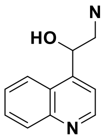 | Mefloquine, Quinine |
| 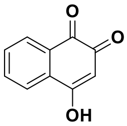 | Atovaquone | 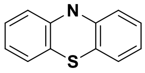 | Methylene Blue |
| 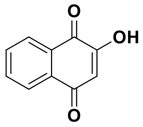 | Atovaquone | 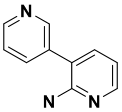 | MMV048 |
| 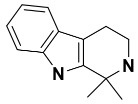 | Cipargamin | 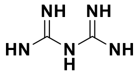 | Proguanil |
| 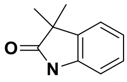 | Cipargamin | 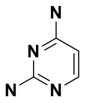 | Pyrimethamine, Diaminopyrimidines |
| 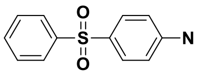 | Dapsone | 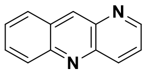 | Pyronaridine |
| 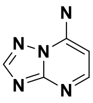 | DSM265 | 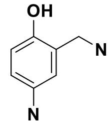 | Pyronaridine side-chain |
| 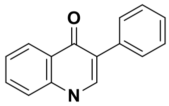 | ELQ-300 | 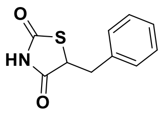 | Rosiglitazone |
| 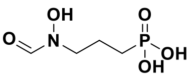 | Fosmidomycin | 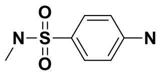 | Sulfadoxine |
| 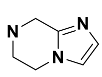 | Ganaplacide | 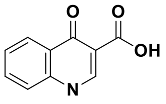 | Temafloxacin |

**Table S4:** Validation of the nanoGlo read-out at PPSC using the MMV Assay Validation Box (63 compounds).

| BATCH_ID | TRIVIAL_NAME | NF175 LDH - $IC_{50}$ - 72h | NF54 nanoGlo - $IC_{50}$ - 72h |
|---|---|---|---|
| MMV000001-17 | Amodiaquine | 0.007 | 0.008 |
| MMV000003-02 | Artemether | 0.007 | 0.008 |
| MMV000004-05 | Dihydroartemisinin | 0.002 | 0.001 |
| MMV000008-29 | Chloroquine | 0.076 | 0.065 |
| MMV000011-03 | Doxycyclin | 1.122 | 1.059 |
| MMV000017-03 | Naphthoquine | 0.292 | 0.367 |
| MMV000019-11 | Artefenomel | 0.004 | 0.003 |
| MMV000022-04 | Piperaquine | 1.821 | 1.479 |
| MMV000024-22 | Pyrimethamine | 0.021 | 20.000 |
| MMV000025-09 | Pyronaridine | 0.031 | 0.045 |
| MMV000028-14 | Trimethoprim | 2.627 | 20.000 |
| MMV000030-02 | Thiostrepton | 0.120 | 0.211 |
| MMV000031-14 | Cycloheximide | 0.164 | 0.145 |
| MMV000037-03 | Fosmidomycin | 6.838 | 5.495 |
| MMV000046-03 | Atovaquone | 0.002 | 0.004 |
| MMV000051-06 | Clindamycin | 0.058 | 0.047 |
| MMV000052-09 | Cycloguanil | 0.007 | 17.179 |
| MMV000055-08 | Sulfadoxine | 20.000 | 20.000 |
| MMV000147-01 | MMV147 | 0.001 | 17.783 |
| MMV001793-06 | Fenretinide | 20.000 | 19.055 |
| MMV002169-13 | Isoniazid | 20.000 | 20.000 |
| MMV002260-06 | Valdecoxib | 20.000 | 20.000 |
| MMV003056-06 | Isoquine | 0.026 | 0.030 |
| MMV003291-01 | Leflunomide | 20.000 | 20.000 |
| MMV004508-06 | 5-Fluorocytosin | 20.000 | 0.001 |
| MMV006169-08 | | 0.288 | 0.351 |
| MMV010036-07 | Panobinostat | 0.009 | 0.010 |
| MMV011565-05 | GNF-Pf-117 | 0.442 | 0.495 |
| MMV016838-02 | | 2.265 | 1.950 |
| MMV018052-10 | MMV052 | 0.005 | 0.003 |
| MMV020752-04 | Pibenzimol | 0.056 | 0.099 |
| MMV034055-05 | ELQ300 | 0.012 | 0.020 |
| MMV1021203-02 | | 1.189 | 2.042 |
| MMV1042856-02 | | 20.000 | 20.000 |
| MMV1042923-02 | | 20.000 | 20.000 |
| MMV1102398-03 | | 20.000 | 20.000 |
| MMV1189010-02 | | 20.000 | 20.000 |
| MMV1191323-02 | | 20.000 | 20.000 |
| MMV1274603-02 | | 20.000 | 20.000 |
| MMV1291222-02 | | 1.567 | 3.388 |
| MMV1402969-02 | | 5.889 | 6.607 |
| MMV1428253-02 | | 2.213 | 3.467 |
| MMV1430193-02 | | 0.417 | 0.724 |
| MMV1432165-02 | | 0.977 | 1.531 |
| MMV1434033-02 | | 12.020 | 8.375 |
| MMV1451822-02 | | 20.000 | 10.839 |
| MMV1533004-02 | | 20.000 | 20.000 |
| MMV1557856-03 | Birinapant | 0.794 | 0.759 |
| MMV1578570-01 | | 0.127 | 0.097 |
| MMV1578889-01 | 680C91 | 20.000 | 13.032 |
| MMV1580173-02 | Trimetrexate | 0.001 | 0.966 |
| MMV1580496-01 | Triapine | 0.367 | 0.417 |
| MMV1634391-01 | MGB-BP-3 | 0.028 | 0.030 |
| MMV390048-02 | MMV048 | 0.034 | 0.052 |
| MMV390482-24 | MMV482 | 0.101 | 0.085 |
| MMV396785-06 | Alexidine | 0.071 | 0.065 |
| MMV637528-05 | Itraconazole | 20.000 | 11.749 |
| MMV642550-03 | | 0.556 | 0.841 |
| MMV665917-05 | | 20.000 | 7.328 |
| MMV669059-10 | DSM421 | 0.080 | 0.150 |
| MMV936405-01 | | 20.000 | 20.000 |
| MMV936413-01 | | 20.000 | 20.000 |
| MMV951872-02 | | 20.000 | 20.000 |

**Table S5:** Parasitological and toxocological properties of the 33 compouds with the Confimed Active status

| MMV-ID | SMILE | PPSC | | | | TCGLS | |
|---|---|---|---|---|---|---|---|
| | | NF54 nanoGlo - $IC_{50}$ (µM) | NF54 LDH - $IC_{50}$ (µM) | Dd2 LDH - $IC_{50}$ (µM) | HepG2 - $IC_{50}$ (µM) | 3D7 LDH - $IC_{50}$ (µM) | HepG2 - $IC_{50}$ (µM) |
| MMV1642139 | Clc1cccc(OC2CCN(C2)C(=O)c3cscn3)c1 | 3.98 | 6.92 | 1.95 | > 10 | 0.580 | > 25 |
| MMV1648843 | CN1[C@@H]2CN(Cc3c(C)onc3C)C[C@@H]2Oc4cc(F)ccc4C1=O | 0.720 | 1.02 | 1.00 | >10 | 0.490 | > 25 |
| MMV1653354 | COc1ccc(cc1OC)C(=O)N2C(CC(c3ccccc3)n4ncnc24)c5ccc(C)cc5 | 0.460 | 1.17 | 0.74 | > 10 | 0.120 | > 25 |
| MMV1656184 | CN1CCC(CC1)(N(C2CCN(Cc3ccccc3)CC2)C(=O)C)C(=O)Nc4ccc5[nH]ccc5c4 | 1.66 | 1.26 | 1.23 | >10 | 1.23 | > 25 |
| MMV1658051 | CC(N)c1ccc(Nc2ncc3cc(ccc3n2)c4ccncc4)cc1 | 1.10 | 2.69 | 1.20 | > 10 | 1.05 | 11.4 |
| MMV1664010 | Cc1ccc2N(CCCc2c1)C(=O)CN3CCC(=CC3)c4cnn(C)c4 | 3.72 | 3.8 | 1.74 | > 10 | 0.777 | > 25 |
| MMV1669145 | Nc1ncc(s1)C(=O)N2CCC3(CCCCCC3)C2 | 0.830 | 0.870 | 0.650 | > 10 | 0.470 | > 25 |
| MMV1677949 | COc1cccc(n1)c2cc(O)cc(CCC3CCCCN3)c2 | 1.23 | 1.82 | 1.55 | > 10 | 1.28 | > 25 |
| MMV1697393 | Cc1cnccc1NCCNCc2cccc(O)c2Cl | 1.12 | 1.48 | 1.23 | > 10 | 0.790 | > 25 |
| MMV1698238 | CC(C)c1ccc(cc1)c2nc(CN3CCCCC3)cc(CN4CCCCC4)c2O | 0.810 | 2.40 | 1.23 | > 10 | 1.13 | > 25 |
| MMV1698806 | Cc1ccc(NC(=O)c2c(C)nc3sc(C(=O)NCc4ccccc4)c(N)c3c2c5ccccc5Cl)c(C)c1 | 0.830 | 1.41 | 1.78 | > 10 | 1.08 | > 25 |
| MMV1700353 | COc1ccc(cc1)c2cc(C(=O)NC3CCN(C3)C4=NNC(=O)C=C4)c(C)n2C | 0.500 | 2.04 | 0.870 | > 10 | 0.480 | > 25 |
| MMV1700355 | COc1ccc(cc1)c2cc(C(=O)NC3CCN(C3)C4=NN(C)C(=O)C=C4)c(C)n2C | 1.05 | 3.63 | 0.790 | > 10 | 0.350 | > 25 |
| MMV1704593 | CCCCN(CC)CCNC(=O)C1CCN(CC1)C2=NS(=O)(=O)C(=C2C)c3ccc(OC)cc3 | 3.02 | 1.20 | 1.32 | > 10 | 0.700 | > 25 |
| MMV1704710 | CCC1=C(c2ccc(C)cc2)S(=O)(=O)N=C1N3CCC(CC3)C(=O)NCCN4CCCCC4C | 2.69 | 1.35 | 1.41 | > 10 | 0.400 | > 25 |
| MMV1722301 | COc1ccc(CC(=O)Nc2cc(C)n(n2)c3cccc(Cl)c3)cc1 | 1.55 | 2.92 | 1.55 | > 10 | 0.560 | > 25 |
| MMV1723349 | CCOc1cccc(c1)c2ccc3nc(c(CCC(=O)NCCN4CCCC4)n3c2)c5cccc(OC)c5 | 0.310 | 0.280 | 1.20 | > 10 | 0.0548 | > 25 |
| MMV1726212 | Oc1ccccc1CCNC2CCc3ccc(F)cc23 | 0.190 | 0.310 | 0.270 | > 10 | 0.270 | > 25 |
| MMV1734041 | Clc1ccc(cc1C#N)S(=O)(=O)NCCN(C2CC2)S(=O)(=O)c3ccc(Cl)c(c3)C#N | 2.04 | 3.31 | 2.00 | > 10 | 1.10 | 13.6 |
| MMV1734413 | NC(=NCc1cccc2cccnc12)N3CCN(CC3)c4nccs4 | 1.70 | 0.450 | 0.500 | > 10 | 0.710 | > 25 |
| MMV1737523 | Clc1cccc(CNCCCNc2ccncc2)c1Cl | 2.00 | 2.00 | 1.55 | > 10 | 1.15 | 16.3 |
| MMV1742686 | CC1(C)OCc2cc(CNc3ccc4OC(=O)Nc4c3)ccc2O1 | 0.550 | 0.760 | 0.660 | > 10 | 0.380 | 18.6 |
| MMV1744039 | Cc1nc(N)nc(C)c1C(=O)Nc2ccc(Nc3ncccn3)cc2 | 1.48 | 1.78 | 1.70 | > 10 | 1.46 | > 25 |
| MMV1747770 | CC(C)C(NCc1ncccc1O)c2oc3ccccc3c2C | 0.280 | 0.830 | 0.600 | > 10 | 0.180 | > 25 |
| MMV1749168 | CN1CCC2(CCN(C2)c3ccnc4ccsc34)C1 | 0.550 | 0.350 | 0.510 | > 10 | 0.500 | 24.9 |
| MMV1752028 | C1CNCC2(C1)CCN(C2)c3ccnc4ccsc34 | 0.480 | 0.390 | 0.400 | >10 | 0.460 | > 25 |
| MMV1752548 | CCn1cnc2c(ncnc12)N3CCCC(C3)c4ccncc4 | 1.00 | 0.790 | 0.710 | > 10 | 0.660 | > 25 |
| MMV1754342 | CC(C)c1ccc2ncnc(NC3CCc4[nH]cnc4C3)c2c1 | 0.590 | 1.00 | 0.590 | > 10 | 0.530 | > 25 |
| MMV1760339 | CCc1nc2c(ccnc2[nH]1)C(=O)N3CCOc4c(C)cc(C)cc34 | 1.41 | 4.27 | 1.29 | > 10 | 0.230 | > 25 |
| MMV1760745 | Fc1ccc(cc1)C(=O)c2ccccc2C(=O)Nc3cnc4ccccc4c3 | 0. 490 | 2.57 | 0.950 | > 10 | 1.39 | > 25 |
| MMV1767993 | Oc1ccc(cc1NC(=O)c2cccc(Oc3cccc(c3)C(F)(F)F)c2)N4CCCS4(=O)=O | 0.350 | 1.10 | 0.540 | > 10 | 0.490 | > 25 |
| MMV1770436 | Cc1ccc(O)c(CNCC2(CCOCC2)c3ccccc3Cl)n1 | 0.210 | 0.580 | 0.440 | > 10 | 0.410 | > 25 |
| MMV1771347 | Cc1cccc(CN(CC2CCCCO2)Cc3cccc(C)n3)n1 | 0.230 | 0.680 | 0.280 | > 10 | 1.32 | > 25 |

**Table S6:** Biological, physiochemical and ADMET properties of the four representative re-confirmed hits.

| | **MMV1722301** | **MMV1669145** | **MMV1747770** |
|---|---|---|---|
| Structure | 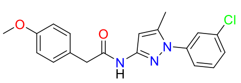 | 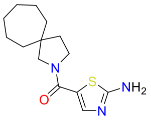 | 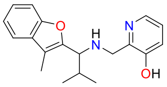 |
| *P.f.* 3D7 LDH $IC_{50}$ (µM) | 0.75 | 0.57 | 0.16 |
| HepG2 $IC_{50}$ (µM) | > 25 | > 25 | > 25 |
| hERG (% inhib. at 1 µM) | < 30 | < 30 | < 30 |
| Solubility (µM) | 13 | 28 | 146 |
| eLogD | 4.11 | 3.07 | 3.19 |
| HLM (µL/min/mg) | 19 | 13 | 464 |
| RHEPs (µL/min/$10^6$ cells) | 22 | 179 | 127 |
| MPO score | 0.15 | 0.47 | 0.53 |

ADMET: absorption, distribution, metabolism, excretion and toxicity; HLM: human liver microsomes; RHEPs: rat hepatocytes; MPO: multi-parameter optimization

**Supplementary methods**

*Metabolic stability study using human liver microsomes* (TCGLS)

A solution of the test compounds in phosphate buffer solution (1 µM) was incubated in pooled human liver microsomes (0.5 mg/mL) for 0, 5, 20, 30, 45 and 60 minutes at 37 °C in the presence and absence of NADPH regeneration system(NRS). The reaction was terminated with the addition of ice-cold acetonitrile containing system suitability standard at designated time points. The sample was centrifuged (4200 rpm) for 20 minutes at 20 °C and the supernatant was half diluted in water and then analyzed by means of LC-MS/MS. % Parent compound remaining, half-life ($T_{1/2}$) and clearance ($CL_{int,app}$) were calculated using standard methodology. The experiment was carried out in duplicate. Verapamil, diltiazem, phenacetin and imipramine were used as reference standards.

*Metabolic stability using cryopreserved rat hepatocytes* (TCGLS)

A solution of the test compound in in Krebs-Henseleit buffer solution (1 µM) was incubated in pooled rat hepatocytes (1×106cells/mL) for 0, 15, 30, 45, 60, 75 and 90 minutes at 37 °C (5% CO2, 95% relative humidity). The reaction was terminated with the addition of ice-cold acetonitrile containing system suitability standard at designated time points. The sample was centrifuged (4200 rpm) for 20 minutes at 20 °C and the supernatant was half diluted in water and then analyzed by means of LC-MS/MS. % Parent compound remaining, half-life ($_{T1/2)}$ and clearance ($CL_{int,app}$) were calculated using standard methodology. The experiment was carried out in duplicate. Diltiazem, 7-ethoxy coumarin, propranolol and midazolam were used as reference standards.

*Solubility in phosphate buffered saline (PBS) pH7.4* (TCGLS)

The solubility assay was performed using a miniaturized shake flask method. A solution of phosphate buffered saline (PBS) and the test compound (200 µM) was incubated at 25 °C with constant shaking (600 rpm) for 2 hours. The samples were filtered using a multiscreen solubility filter plate. The filtrate was half diluted in acetonitrile. A five-point linearity curve was prepared in PBS:Acetonitrile (1:1, v/v) at 200, 150, 75, 25 and 2.5 µM. Blank, linearity and test samples (n = 2) were transferred to a UV readable plate and the plate was scanned for absorbance. Best fit calibration curves were constructed using the calibration standards and used to determine the test sample solubility. The experiment was carried out in duplicate. Diethylstilbestrol, haloperidol and sodium diclofenac were used as reference standards.

*LogD at pH7.4* (TCGLS)

The LogD pH7.4 assay was performed using a miniaturized shake flask method. A solution of a pre-saturated mixture of 1-octanol and phosphate buffered saline (PBS) (1:1, v/v) and the test compound (75 µM) was incubated at 25 °C with constant shaking (850 rpm) for 2 hours with The organic and aqueous phases were separated, and samples of each phase transferred to plate for dilution. The organic phase was diluted to 1000-fold and the aqueous phase was diluted 20-fold. The samples were quantitated using LC-MS/MS. The experiment was carried out in duplicate. Propranolol, amitriptyline and midazolam were used as reference standards.

*Measurement of hERG binding*

Affinity for the hERG (human ether-a-go-go-related-gene) ion channel was evaluated using a Predictor™ hERG Fluorescence Polarization Assay kit (Invitrogen, Catalog no: PV5365). The test compound (1 µM) was incubated at ambient temperature for 4 hours with hERG membrane and red fluorescent hERG channel ligand (provided in the kit). A DMSO concentration of 1% was maintained in all wells. Fluorescence polarization was measured at an emission wavelength of 595 nm with the excitation of 531 nm using a microplate reader (Envision, Perkin Elmer). E-4031 was used as the reference inhibitor. Inhibition for test compounds was calculated considering the mP values of E-4031 (30 µM) as 100% inhibition and vehicle control as 0% inhibition
